# CD39 defines a cytotoxic tumor-associated NK cell state responsive to NKG2A blockade in lung cancer

**DOI:** 10.1101/2025.08.26.672316

**Authors:** Clara Serger, Lucas Rebuffet, Michael T. Sandholzer, Wiebke Rackwitz, Irene Fusi, Petra Herzig, Annalea Brüggemann, Carina A. Pawlow, Evgeny Chichelnitskiy, Daniel Hajnal, Nicole Oelgarth, Sarp Uzun, Christoph Schultheiß, Andreas Zingg, Sofia Tundo, Thuy T. Luu, Aljaz Hojski, Didier Lardinois, Lavinia Neubert, Mascha Binder, Kirsten Mertz, Marcel P. Trefny, Nicole Kirchhammer, Marina Natoli, Matthias S. Matter, Heinz Läubli, Christine Falk, Karin Schaeuble, Eric Vivier, Andrea Romagnani, Alfred Zippelius

## Abstract

Natural killer (NK) cell-targeting immunotherapies are emerging, yet the differentiation and functional states of tumor-infiltrating NK cells remain poorly understood. Using matched single-nucleus RNA and ATAC sequencing of non-small cell lung cancer (NSCLC) specimens, we resolved the transcriptional and epigenetic landscape of intratumoral NK cells. We identified two tumor-associated NK (taNK) cell subsets marked by *ITGAE* (CD103) and *ITGA1* (CD49a) that display features of tissue residency and dysfunction while preserving cytotoxic function. Trajectory and regulon analyses revealed an inflammation-driven transition from early *GZMK*⁺ NKs toward an *ENTPD1*^+^ (CD39⁺) effector state characterized by interferon-stimulated gene (ISG) programs. Functional profiling established CD39⁺ taNK as the dominant cytotoxic NK cell population with superior killing capacity that is further potentiated by NKG2A blockade. This study offers mechanistic insights into NK cell differentiation in NSCLC and establishes CD39⁺ taNK cells as a targetable effector population for immunotherapy.

**One sentence summary:** Multiomic analysis identifies CD39^+^ tumor-associated NK cells as a cytotoxic effector state in NSCLC responsive to NKG2A blockade.

## INTRODUCTION

Lung cancer is the leading cause of cancer-related mortality worldwide, with NSCLC accounting for approximately 85% of cases (*1*). Although T cell-based immunotherapies have improved survival rates in NSCLC, durable responses are limited to a subset of patients, emphasizing the need for improved therapeutic approaches (*2*).

Besides T cells, NK cells have gained attention as promising immunotherapeutic target. Their intratumoral abundance has been linked to improved overall survival in various cancer types (*3–6*) and is predictive of response to immune checkpoint blockade (ICB) in NSCLC (*7, 8*). These findings have driven the development of numerous NK cell-directed therapies, many of which are currently being evaluated in clinical trials (*9*). The therapeutic efficacy of NK cell-based interventions in solid tumors has been limited by multiple challenges, including NK cell dysfunction, poor tumor homing, and immunosuppressive factors such as Transforming Growth Factor β (TGF-β) (*10*). A deeper understanding of intratumoral NK cells and their transcriptional dynamics within the tumor microenvironment (TME) is essential to advance the development of more effective NK cell-targeting approaches.

In peripheral blood, human NK cells can be categorized into three main subsets: NK1, NK2, and NK3 (*11*). NK1 cells (CD56^dim^CD16^+^) exhibit high cytotoxic potential, whereas NK2 cells (CD56^bright^CD16^−^) are primarily involved in immunoregulatory functions through cytokine secretion (*12*). NK3 cells, distinguished by high levels of NKG2C, reflect a mature phenotype, resembling NKG2C^+^ memory NK cells observed following HCMV infection (*11, 13*). In contrast to circulating subsets, tissue-resident NK cells exhibit unique transcriptional and functional profiles that vary across different organs, shaped by their local microenvironment. These cells are typically less mature and share similarities with CD56^bright^ NK cells (*14*). Overall, NK cells are heterogenous, and their differentiation, activation state, and functional capacity are shaped by their localization and surroundings.

NK cells in solid tumors often acquire a dysfunctional phenotype, characterized by increased expression of inhibitory receptors, coupled with down-regulation of activating receptors, compromising their tumor-killing capacity (*15–20*). Recent studies in human lung tumors have identified NK cells expressing markers associated with tissue-residency (CD103, CD49a) and dysfunction (*21, 22*). Similar observations have been made in colorectal carcinoma (CRC), where CD49a^+^ NK cell frequencies expand with tumor progression (*23*). Additionally, single cell studies of intratumoral NK cells across multiple cancer types have revealed the consistent emergence of dysfunctional, stressed, and tissue-resident phenotypes upon tumor infiltration(*18, 24–26*). Despite these advances, the precise intratumoral NK cell states, along with their underlying gene regulatory networks (GRNs) and environmental cues that govern their differentiation, remain poorly understood. Gaining a deeper understanding of these mechanisms is essential for elucidating the origins of NK cell dysfunction and guiding the development of more effective NK cell-based immunotherapies.

To address this gap, we applied integrated single-nucleus RNA and ATAC sequencing (snRNA-seq and snATAC-seq) to map the transcriptional landscapes and GRNs of NK cells in treatment-naïve NSCLC. Our analysis revealed two distinct tumor-associated (taNK) states, marked by the expression of *ITGAE* (CD103) and *ITGA1* (CD49a), that, while exhibiting features of tissue residency and dysfunction, maintain a pronounced effector profile potentially shaped by inflammatory signaling. Furthermore, we identified a CD39⁺ subset within the CD103⁺CD49a⁺ taNK population that represents a highly cytotoxic state whose anti-tumor activity is further augmented by NKG2A blockade. Together, these findings provide a refined framework of NK cell heterogeneity in NSCLC and establish CD39⁺ taNK cells as an attractive target for immunotherapeutic intervention.

## RESULTS

### NK cells from human lung cancer exhibit substantial heterogeneity

To investigate NK cell heterogeneity in NSCLC and the underlying GRNs shaping their functional states, we performed paired snRNA-seq and snATAC-seq using the 10x Genomics Multiome platform on CD56^+^ immune cells isolated from tumor resections of 11 treatment-naïve patients diagnosed with either lung adenocarcinoma (LUAD, n = 7) or lung squamous cell carcinoma (LUSC, n = 4) (Fig. 1A and fig. S1A, table S1). After quality control and filtering, we recovered 57,854 cells, comprising T cells, NK cells, and innate lymphoid cells (ILCs), with smaller populations of B cells and monocytes. NK cells were computationally enriched by removing T cells, monocytes, B cells, and regulatory T cells. Subsequent analyses were performed following standard Seurat (*27*) and Signac(*28*) workflows (fig. S1, B to D).

**Fig. 1.**
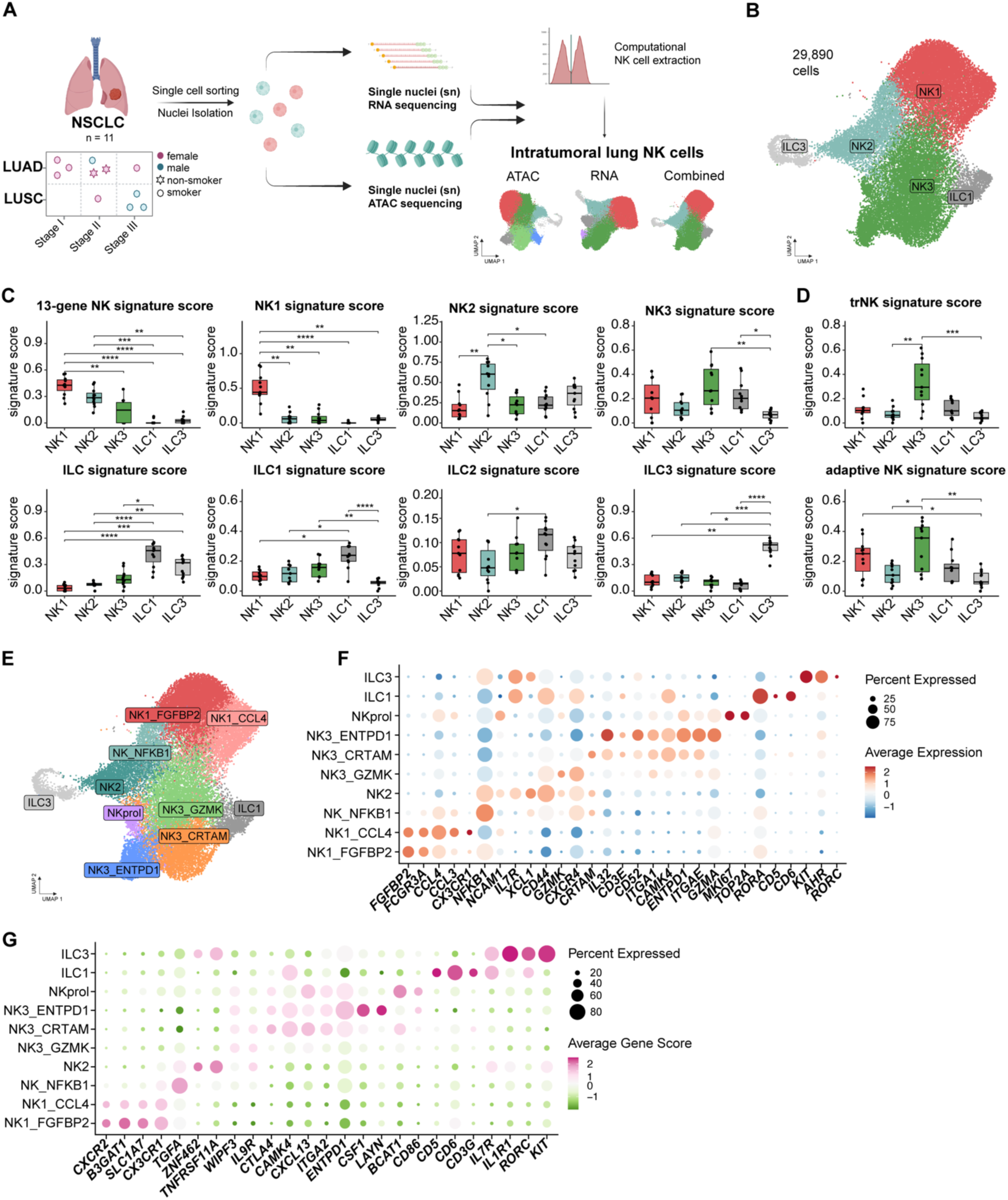
Natural Killer (NK) cells from human lung cancer exhibit substantial heterogeneity. **(A)** Study overview and patient classification. Single cells were sorted from resected lung tumor tissues (n = 11), nuclei were isolated to perform single-nucleus RNA (snRNA) and single-nucleus ATAC (snATAC) sequencing using the 10x Multiome kit. Natural killer (NK) cells were extracted, ATAC and RNA Uniform Manifold Approximations and Projections (UMAPs) were clustered and integrated, leading to a final combined UMAP (Graphics created using BioRender.com (*142*)). **(B)** UMAP representation of combined ATAC and RNA information using WNN integration. Coarse clustering is shown. **(C** and **D)** Box and Whisker plots representing U cell scores for published NK (*11–13, 31*) and ILC (*30*) signatures across coarse NK cell clusters. Individual data points correspond to median scores per sample . **(E)** UMAP showing identified fine clusters of NK cells. **(F** and **G)** Dotplot showing column-scaled gene expression and gene scores of indicated genes. Data are based on single-cell analysis from n = 11 patient samples. Statistics in (C) and (D): non-parametric Kruskal-Wallis test with post-hoc Dunn’s test, adjusted *P* value. (*\*P < 0.05, **P < 0.01, ***P < 0.001, ****P < 0.0001*)

To integrate snRNA-seq and snATAC-seq modalities, we employed weighted nearest neighbor (WNN) analysis (*29*), recovering 29,890 high-quality cells and identifying six distinct clusters (fig. S1E). Cluster 5, present in only one patient and characterized by a pronounced stress-associated gene expression profile (*26*), was excluded due to its patient-specific nature (fig. S1, E to G). The remaining five clusters were annotated based on the expression of key marker genes and chromatin accessibility in proximity to indicated genes (Fig. 1B and fig. S2A). Annotations were validated using publicly available NK and ILC signatures (*11, 12, 30*), identifying profiles resembling NK1, NK2, and NK3 cells in NSCLC lesions (Fig. 1C and fig. S2B, data file S1). NK1 cells were characterized by *NKG7* and *FCGR3A* expression and chromatin accessibility near *B3GAT1* and *CX3CR1*. NK2 cells were defined by elevated levels of *XCL1* and *CD44*, with accessible chromatin near *ZNF462* and *ZNF667*. NK3 cells were marked by *CD52* and *IL32* expression and chromatin accessibility near *CTLA4* and *ITGA2*. Additionally, we observed clusters resembling ILC1s (*RORA, CD6*) and ILC3s (*KIT, RORC*) (fig. S2, A and B). Further comparison with published NK cell signatures revealed that NK3 cells were strongly enriched for a lung tissue-resident NK signature, indicating their potential localization within the tissue (Fig. 1D, data file S1) (*31*). Consistent with previous studies (*11*), the NK3 population also displayed features of adaptive NK cells(*13*), supporting the presence of an adaptive-like NK cell subset in human lung cancer.

To further resolve NK cell diversity, we increased the clustering resolution and performed differential gene expression (DGE) and gene activity analyses (Fig. 1E and fig. S2C, data file S2). NK1 cells segregated into two subclusters; NK1_CCL4, characterized by high expression of chemokines and chemokine receptor *such as CCL4, CCL3, and CX3CR1*; and NK1_FGFBP2, marked by elevated expression of *FGFBP2*. The NK3 population divided into three transcriptionally distinct sub-clusters. NK3_GZMK cells expressed genes commonly associated with CD56^bright^ NK cells (*11, 12*), such as *CD44* and *GZMK*, while NK3_CRTAM and NK3_ENTPD1 were enriched for transcripts linked to tissue-residency (*ITGAE, ITGA1*), adaptive NK cells (*CD3E, IL32*) (*11, 13*), and cytotoxicity (*GZMA*). These subsets also showed chromatin accessibility at loci associated with dysfunction and tumor infiltration, including *CTLA4, CXCL13,* and *LAYN* (*32*) (Fig. 1, F and G). NK_NFKB1 and the proliferating NK cluster (NKprol) shared overlapping features with multiple NK subtypes (Fig. 1, F and G). Of note, no association was observed between NK subset composition and NSCLC histology or tumor stage, likely due to limited sample size (fig. S2, D and E).

In summary, our integrated single nucleus analysis reveals a complex NK cell landscape in NSCLC, consisting of multiple distinct NK1, NK2, and NK3 subpopulations. In addition, the NK3 subset identified here not only mirrors circulating NK3 cells but also exhibits unique features of tissue residency and adaptive NK cells.

### Cross-modality analysis suggests distinct epigenetic origins

To investigate potential differences between the epigenetic and transcriptional cell states that could indicate distinct lineage origins, we performed a comparative cross-modality analysis. RNA and ATAC datasets were clustered independently, and cluster assignments were correlated with integrated WNN clusters to identify modality-specific features (fig. S3, A to C). We identified two distinct chromatin states, ATAC Cluster a2 and a6, which mapped primarily to a single transcriptional state, rNK3_ENTPD1, with a smaller fraction aligning to rNK3_GZMK (fig. S3, D and E). The subsequent WNN integration resolved these epigenetic differences: NK3_CRTAM corresponded primarily to ATAC Cluster a2, whereas NK3_ENTPD1 aligned with ATAC Cluster a6 (fig. S3, D and F). In addition, both WNN subclusters remained transcriptionally embedded within the broader rNK3_ENTPD1 RNA cluster (fig. S3, D and G).

In addition, the NK_NFKB1 cluster maps across multiple RNA lineages (NK1 and NK2) and ATAC clusters (fig. S3D). Similarity analysis (fig. S3, F and G) shows that the NK_NFKB1 cluster exhibits high transcriptional and gene activity correlations with both, NK1 and NK2, supporting its identity as a cluster of combined cell states.

Collectively, multimodal integration is required to resolve epigenetically distinct fates, such as NK3_CRTAM and NK3_ENTPD1, which represent distinct lineages that converge transcriptionally, while retaining their ancestral chromatin marks.

### Diverse GRNs and transcriptional profiles reveal functional specialization of NK cell subsets in NSCLC

Recognizing that marker gene expression alone provides an incomplete view of cell states, we aimed to reconstruct the transcriptional programs and GRNs that shape NK cell fate in NSCLC. We first assessed transcription factor (TF) motif accessibility, identifying binding motifs for well-established NK cell TFs, including *EOMES* and *TBX21*, across major NK cell subsets (fig. S3H). To systematically map transcriptional regulation, we next applied SCENIC+, a multiomics framework that integrates TF expression, enhancer accessibility, and target gene co-expression, to infer enhancer-driven GRNs (*33*). This approach enabled us to define active regulatory modules across NK cell subsets in lung tumors.

The analysis confirmed known regulatory modules, including strong *RORC* and *AHR* regulon activity in ILC3s, high E2F family regulon activity in proliferating NK cells, and enrichment of NFKB1-associated regulons (*34, 35*) in the NK_NFKB1 cluster (Fig. 2A). In addition, ILC1s displayed strong *RORA* regulon activity, reinforcing its utility as a distinguishing candidate GRN between NK cells and ILC1s in NSCLC (*36, 37*). Within the NK1 clusters, as previously reported (*38*), we observed enrichment of regulons associated with NK cell cytotoxicity, effector function and maturation, including *IRF1* (*39, 40*) and *PRDM1* (*41*). The NK2 subset showed enhanced activity of *TCF7* (*38*) and *BACH2* (*42*) regulons, both linked to immature NK cells. In contrast, NK3 clusters were marked by strong activity of *PRDM1*, a regulon implicated in NK maturation (*41*) and the NK3 phenotype (*11*), and *RUNX2*, previously associated with tissue residency (*43*).

**Fig. 2.**
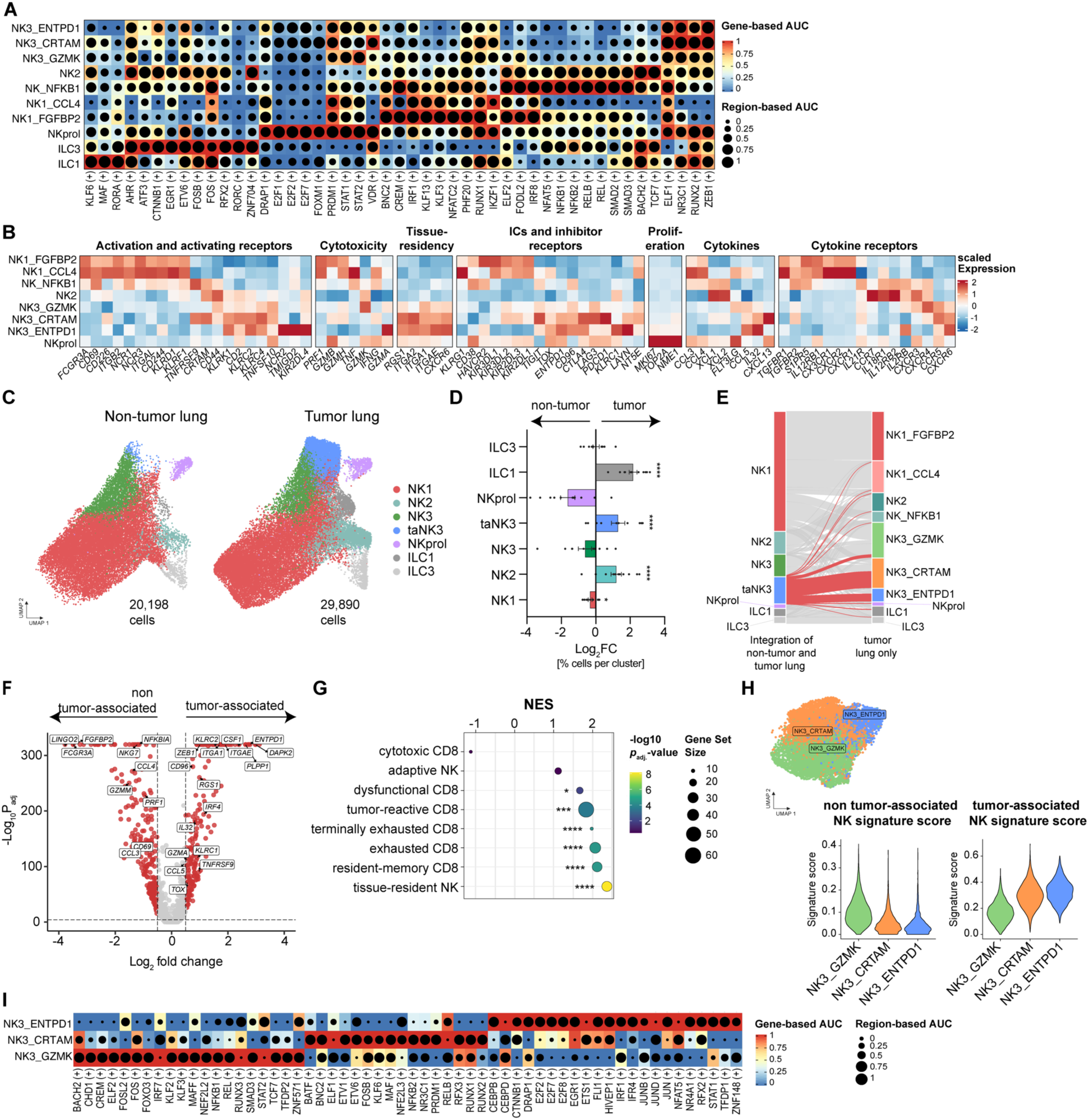
NK cells exhibiting a tissue-resident and dysfunctional profile are enriched in tumor tissue. (A and. **B)** Heatmap visualizing the differential activity of (A) enhancer-driven regulons, indicated by (+), and (B) gene expression of indicated genes in NK cell clusters (ICs = immune checkpoints). **(C)** Integrated UMAP representation of non-tumor lung (Sikkema *et al.* (*62*), Bischoff *et al.* (*63*)) and tumor lung NK cells split by tissue origin (ta = tumor-associated). **(D)** Bar graph depicting mean ± SEM of log2 fold change (Log2FC) of cell frequencies in each cluster from individual lung tumor samples, compared to the average frequencies of the corresponding clusters in non-tumor lung tissue. (tumor lung: n=11, non-tumor lung: n=50) **(E)** Alluvial plot showing the distribution of integrated non-tumor and tumor lung NK cells across clusters identified in the tumor lung NK cell UMAP. **(F)** Volcano plot depicting differentially expressed genes (DEGs) between tumor-associated and non tumor-associated NK cells. **(G)** Dot plot showing normalized gene set enrichment analysis (GSEA) scores (NES) of published signatures (*13, 31, 54, 65–66, 119*) in tumor-associated NK (taNK) cells signature. **(H)** UMAP visualization of re-clustered NK3 subsets (left) alongside violin and feature plots (right) showing the enrichment of the taNK signature and the non-taNK signature (top 30 genes) within these NK3 clusters. **(I)** Heatmap displaying the differential activity of enhancer-driven regulons, indicated by (+), across each NK3 cluster. Data were analyzed by (D) Wilcoxon signed-rank test on raw frequency values, (F) using the MAST hurdle model and *P*-values were calculated using a likelihood ratio test and adjusted for multiple comparisons using Bonferroni correction, (G) permutation-based testing with adjustment for multiple comparisons using ClusterProfiler. *P < 0.05, **P < 0.01, ***P < 0.001, ****P < 0.0001

To further elucidate the functional potential of these NK cell subsets, we categorized genes based on their roles in activation, cytotoxicity, tissue-residency, immune checkpoints and inhibitory receptors, proliferation, cytokines, and cytokine receptors (Fig. 2B). NK1 and NK2 clusters mirrored transcriptional profiles previously characterized in peripheral NK cells: NK1 cells expressed a broad array of activating and cytolytic genes, as well as genes associated with tissue migration (*GZMB, FCGR2A, S1PR5*, *CX3CR1*), whereas NK2 cells expressed cytokine-producing genes (*XCL1, XCL2*) (*11, 44, 45*). In contrast, NK3 cells exhibited distinct activation-related profiles and a strong enrichment for tissue residency–associated genes such as *ITGA1*, *ITGAE*, *CXCR6*. Within NK3 subsets, NK3_CRTAM cells expressed *CRTAM, CD44,* as well as elevated levels of *KLRC2* (NKG2C) and *TNFRSF9,* genes linked to adaptive NK cells (*13*) and activation (*46*), respectively. NK3_ENTPD1 cells expressed high levels of *NCR2* and *TMIGD2*, a CD28 family member known to promote NK cell proliferation and cytokine production (*47, 48*). While cytotoxicity-related gene expression appeared reduced in NK3 cells, NK3_CRTAM cells retained *IFNG* expression, and NK3_ENTPD1 cells expressed *GZMA*, indicating effector potential. Both NK3 subsets also exhibited transcriptional hallmarks of dysfunction and chronic T cell stimulation (*32*), including classical immune checkpoints such as *TIGIT*, *CTLA4*, and *LAG3*.

Additionally, NK3_ENTPD1 cells expressed high levels of *KLRC1* (NKG2A), a marker associated with dysfunctional intratumoral NK cells (*49*), and *LAYN*, a gene frequently associated with CD8^+^ T cell dysfunction (*50*) (Fig. 2B).

With regard to cytokines and chemokines, NK3 cells expressed *CCL5*, which supports recruitment of CD8^+^ T cells and DCs (*51, 52*), and *IL32*, a cytokine involved in anti-tumor immunity (*53*). Aditionally, NK3_CRTAM cells showed high expression of *CXCL13*, a chemokine enriched in tumor-reactive T cells (*54, 55*), critical for tertiary lymphoid structure (TLS) formation (*56*), and predictive for response to ICB (*57, 58*). Elevated expression of *CXCR6*, a molecule crucial for memory NK cell persistence (*59*) and T cell-mediated tumor control (*60*), was also observed, supporting its role in tissue retention of NK cells in NSCLC (*61*) (Fig. 2B).

In conclusion, NK cells in NSCLC demonstrate a broad spectrum of GRNs and transcriptional programs associated with distinct functional identities. These include conventional cytotoxic NK1 and NK2 subsets, as well as functionally diverse NK3 populations with tissue-resident, adaptive, and dysfunctional features.

### NK cells exhibiting a tissue-resident and dysfunctional profile are enriched in tumor tissue

To identify NK cell subtypes specifically associated with the TME, we integrated our snRNA-seq dataset from NSCLC with two publicly available datasets of non-tumor lung NK cells (*62, 63*). Using the standard Seurat workflow, we obtained a combined dataset of 50,088 high-quality NK cells, comprising 20,198 from non-tumor tissues and 29,890 from tumors (Fig. 2C and fig. S4A). Clustering and UMAP visualization revealed seven NK cell subsets, which were annotated based on DGEs and published NK and ILC transcriptional signatures (*11, 12, 30*) (fig. S4, B and C, data file S1). Among these, a distinct NK3 cluster was exclusively present in tumor samples and characterized by high expression of *ITGAE, ITGA1,* and *ENTPD1* (Fig. 2, C and D and fig. S4B and S4D). We designated this subset as tumor-associated NK3 (taNK3). Consistent with previous studies (*15, 17, 64*), we observed a general shift in NK cell composition within tumors, including a reduction of NK1 cells, and a relative enrichment of NK2 and ILC1 cells (Fig. 2D and fig. S4D). The identified taNK3 cluster showed strong overlap with the previously identified NK3_CRTAM and NK3_ENTPD1 clusters (Fig. 2E), suggesting that these two subsets are enriched or expanded in the TME. These two subclusters where thus collectively designated as taNK, and the remaining NK cell clusters were categorized as non-tumor-associated NK cells (non-taNK) (fig. S4E).

We next compared taNK cells (NK3_CRTAM and NK3_ENTPD1) to non-taNK cells from our dataset to identify transcriptional features distinguishing this tumor-associated state. DGE analysis revealed that taNK cells are marked by elevated expression of genes linked to tissue-residency (*ITGAE, ITGA1, ENTPD1*), dysfunction (*ENTPD1, CD96, KLRC1*), cytotoxicity and inflammation (*IL32, CCL5, GZMA*), and adaptive NK cells (*KLRC2*, *IRF4*) (Fig. 2F, data file S2). Gene signature enrichment analysis (GSEA) further confirmed that taNK cells are significantly enriched for a lung tissue-residency signature (*31*), as well as gene programs conventionally associated with dysfunctional and exhausted CD8^+^ T cells (*65, 66*). In addition, taNK cells strongly enriched for tumor-reactive and tissue-resident memory (T_RM_) CD8^+^ signatures, the latter being associated with favorable prognosis and improved responses to ICB across multiple cancer types (*67–70*) (Fig. 2G). We also observed a positive, albeit non-significant, enrichment for adaptive NK cells (*13*), whereas a cytotoxic gene signature was negatively enriched (*65*).

As these analyses compared taNKs against all NK subset, we next aimed to elucidate NK3-specific transcriptional changes and regulon activity. Subclustering of NK3 cells confirmed the three previously identified NK3 clusters: NK3_GZMK, NK3_CRTAM, and NK3_ENTPD1 (Fig. 2H, left panel). Signature enrichment further validated NK3_CRTAM and NK3_ENTPD1 as tumor-associated subsets, both showing high enrichment for the taNK signature. Conversely, NK3_GZMK was enriched for the non-taNK signature, supporting its identity as a non-tumor lung NK3 population (Fig. 2H, right panel; fig S4F, left panel). The intra-NK3 differential expression profile closely mirrored the global taNK signature, underscoring their dysfunctional and tissue-resident phenotype within the TME (fig. S4F, right panel; fig. S4G). To define taNK-specific GRNs, we conducted a SCENIC+ regulon analysis (Fig. 2I), revealing distinct regulatory programs across NK3 subsets. Non-tumor NK3_GZMK were enriched for regulons associated with immaturity (*BACH2* (*42*), *TCF7* (*38*)) and homeostatic maintenance (*KLF2* (*71*)). In contrast, the taNK clusters exhibited GRNs consistent with functional adaptation to the TME: NK3_CRTAM cells were characterized by programs associated with immune suppression (*NR3C1* (*72*)), maturation (*PRDM1* (*41*)), and tissue residency (*RUNX2* (*43*)), whereas N3_ENTPD1 displayed GRNs associated with proliferation (E2Fs (*35*)), dysfunction (*NR4A1* (*73, 74*)), and adaptive responses (*IRF4* (*75–77*)). Importantly, both taNK clusters showed increased activity of AP1 family regulons (*BATF, FOSB, JUN, JUNB, JUND*), previously implicated in adaptive NK cells (*13*).

Lastly, to dissect the transcriptional heterogeneity within the taNK population, we directly compared NK3_CRTAM and NK3_ENTPD1. NK3_CRTAM cells showed elevated expression of *CRTAM* and *GZMH*, while NK3_ENTPD1 cells expressed higher levels of *GZMA* and *FCER1G* (fig. S4H, data file S2). Based on the elevated expression of *BCL11B* (*78*) within the NK3_CRTAM cluster and established evidence of adaptive-like NK cells within the lung environment (*79*), we next investigated whether one of taNK subsets may represent an adaptive-like population. Using a validated human adaptive NK cell dataset, we performed signature enrichment analysis (*13*). Although not statistically significant, this analysis revealed a trend toward higher enrichment of adaptive features in NK3_CRTAM compared to NK3_ENTPD1, suggesting a closer transcriptional alignment with adaptive NK cells (fig. S4I).

In summary, we identified NK cell subsets uniquely associated with the TME with features of NK3 cells, characterized by tissue-residency, and dysfunctionality, closely resembling tumor-reactive CD8^+^ T_RM_ cells, positioning them as potential players in the immune response to lung cancer.

### Multiplexed profiling validates the identity of tumor-associated NK cells

To confirm the transcriptional profile of taNK cells at the protein level, we performed flow cytometry on NK cells isolated from an independent set of NSCLC tumor samples (n = 15), adjacent non-tumor lung tissue (n = 13), and peripheral blood (n = 12) (table S2). NK cells were defined as CD45^+^CD14^-^CD19^-^CD3^-^CD56^+^. As CD103 (*ITGAE*) and CD49a (*ITGA1*) were among the most DEGs distinguishing taNK cells from non-taNK cells, we used those as markers to identify taNK cells (Fig. 2F and fig. S5A). This analysis confirmed a significant enrichment of CD103^+^CD49a^+^ NK cells in tumor lesions compared to adjacent non-tumor lung tissue. In addition, these cells were absent in peripheral blood (Fig. 3, A and B and fig. S5B), underscoring their tumor-restricted localization in NSCLC. Consistent with our transcriptional analyses, taNK cells exhibited elevated protein levels of CCL5, granzyme A (*GZMA*), NKG2A (*KLRC1*), NKG2C (*KLRC2*), CD39 (*ENTPD1*), and CTLA4, alongside reduced expression of granzyme B (*GZMB*), and CD16 (*FCGR3A*) compared to CD103^-^CD49a^-^ NK cells (Fig. 3, C and D and fig. S5C).

**Fig. 3.**
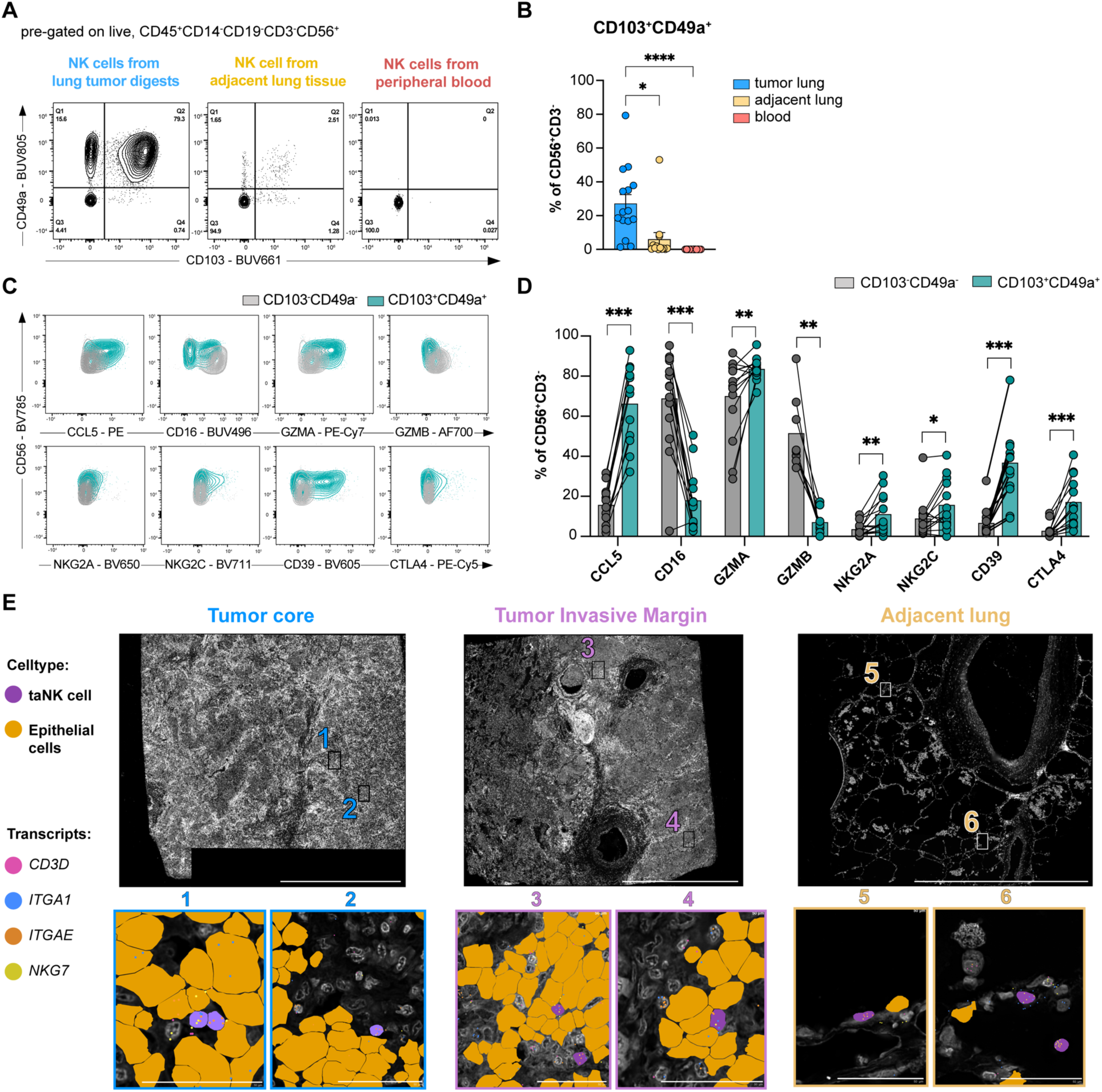
Multiplexed profiling validates the identity of tumor-associated NK cells. **(A)** Representative contour plots showing CD103 and CD49a protein expression in NK cells from lung tumor, adjacent lung and periphery. **(B)** Frequency (mean ± SEM) of CD103^+^CD49a^+^ NK cells in tumor lung and adjacent lung quantified using flow cytometry. (tumor lung: n = 15, adjacent lung: n = 13, blood: n = 12) **(C)** Representative contour plots of indicated proteins in NK cells from tumor digests comparing CD103^+^CD49a^+^ NK cells (petrol) and CD103^-^CD49a^-^ NK cells (grey). **(D)** Barplots showing the mean frequencies of indicated proteins comparing CD103^+^CD49a^+^ NK cells (petrol) and CD103^-^CD49a^-^ NK cells (grey) from tumor lung. (n = 14). **(E)** Representative 10X Xenium images of tumor core, invasive margin, and adjacent lung tissue from one lung adenocarcinoma (LUAD) patient (LuT-722). Panels show DAPI staining (grey) with cell-type masks indicating tumor-associated NK (taNK) cells (purple) and epithelial cells (yellow). Zoomed-in regions (1–6) illustrate the co-localization of *ITGAE*, *ITGA1*, and *NKG7* transcripts within taNK cells. *CD3D* (pink) was utilized to verify the exclusion of T cell contamination. Scale bars: Overview: 2 mm; Zoom-ins: 50 μm. Each dot in (B) and (D) represents an individual sample, and data were analyzed using (B) Kruskal-Wallis test with post-hoc Dunn’s multiple comparison and (D) paired Wilcoxon signed-rank. *\*P < 0.05, **P < 0.01, ***P < 0.001, ****P < 0.0001*

To further investigate the co-expression pattern of *ITGAE* (CD103) and *ITGA1* (CD49a) *in situ* of taNK cells, we performed 10X Genomics Xenium spatial transcriptomics on a representative LUAD patient (LuT-722) comparing tumor core, invasive margin, and adjacent non-tumor lung. Following initial cell segmentation and unsupervised clustering, we refined the NK cell population by excluding *CD3D*-expressing T cells to ensure a high-purity NK cluster for further analysis (fig. S6, A to D). We then applied our discovery taNK signature from the Multiome data to these purified NK cells, utilizing a signature enrichment threshold to identify the taNK subset *in situ* (fig. S6, E and F). Projecting these cells back onto the tissue sections confirmed the coexpression of *ITGAE* (CD103) and *ITGA1* (CD49a) transcripts within individual taNK cells localized in close proximity to epithelial cells, likely tumor cells (Fig. 3E). In addition, in this LUAD patient we observed a numerically higher taNK density within the tumor core and the invasive margin compared to adjacent lung tissue (fig. S6G).

Thus, in accordance with our flow cytometry and Multiome data, this data supports the enriched presence of the taNK cell state within the TME.

### Functional profiling reveals a robust effector identity of taNK cells

Given the enrichment for inhibitory receptors and effector molecules, we assessed the functional responsiveness of taNK cells from lung tumor lesions (n = 8) using either a cytokine cocktail (IL-12, IL-15, and IL-18) or coculture with irradiated K562 cells (Fig. 4A, table S2). Even without stimulation, CD103^+^CD49a^+^ taNK displayed higher Ki67 expression compared to CD103^-^CD49a^-^non-taNK, consistent with elevated *MKI67* transcripts, indicating an active cell cycle (Fig. 4B and Fig. 2B). While taNK cells initially showed lower or comparable expression of IFNγ, CD107a, and granzyme B compared to non-taNK cells, they matched or exceeded these levels following 24 hours of cytokine stimulation (fig. S7A). Additional analysis revealed a significantly greater induction of CD107a, granzyme B, and Ki67 in taNK cells, suggesting that these cells are primed to respond to inflammatory cytokines (Fig. 4C). However, in the K562 co-culture, only CD107a showed a significantly increased fold-change in taNK cells, indicating that direct tumor cell stimulation fails to reactivate taNK cells (fig. S7, B and C).

**Fig. 4:**
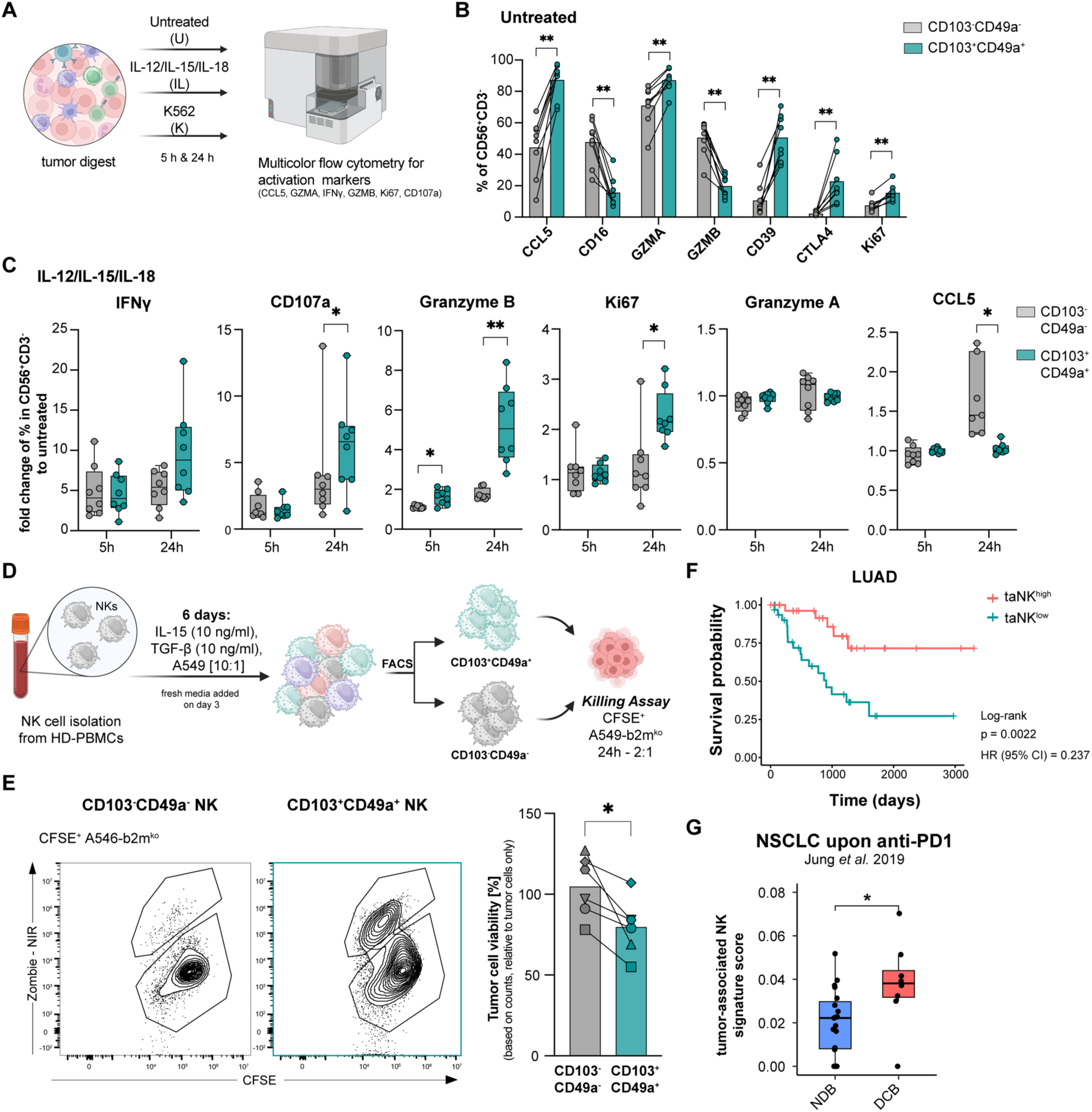
Functional profiling reveals a robust effector identity and enhanced cytotoxicity of tumor-associated NK cells. **(A)** Schematic respresentation of functional assays (Graphics created with BioRender.com (*143*)). **(B)** Barplots showing mean frequencies of indicated protein expressions comparing CD103^+^CD49^+^with CD103^-^CD49^-^ NK cells from lung tumor digests after 5 hours incubation, kept untreated (n = 8). **(C)** Fold change in percentage of indicated markers in CD103^+^CD49a^+^ (petrol) and CD103^-^CD49a^-^NK cells (grey), relative to untreated, from lung tumors upon IL-12/IL-15/IL-18 for 5 or 24 hours. (5 h: n = 8, 24 h: n = 8). **(D)** Schematic representation of killing assays (Graphics created with BioRender.com (*144*)). **(E)** Representative contour plots of ZombieNIR^+^ CFSE^+^ A549-b2m^ko^ following 24 hours co-culture with either sorted CD103^-^CD49a^-^ or CD103^+^CD49a^+^ NK cells from the *in vitro* taNK platform (left). Tumor cell viability was determined by absolute counts and normalized to tumor cell only, comparing CD103^-^CD49a^-^with CD103^+^CD49a^+^ NK cells from the *in vitro* taNK platform (n = 6). **(F)** Kaplan-Maier curve showing the effect of tumor-associated NK cells on overall survival in lung adenocarcinoma (LUAD), (ta = tumor-associated, taNK^high^: top 25%, taNK^low^: bottom 25%). **(G)** Boxplots showing UCell signature score for tumor-associated NK cells in non-small cell lung cancer (NSCLC) upon anti-PD1 treatment grouped by response (NDB: no durable benefit, n = 17; DCB: durable clinical benefit, n = 8). Data were analyzed using [(B), (C) and (E)] paired Wilcoxon signed-rank test, (F) log-rank test, (G) Wilcoxon signed-rank test. *\*P < 0.05, **P < 0.01, ***P < 0.001, ****P < 0.0001*

The limited yield and reduced viability of NK cells isolated from tumor lesions precluded obtaining sufficient cell numbers for robust cytotoxicity assays. To overcome this constraint, we established an in vitro system that models the taNK phenotype using PBMC-derived NK cells from healthy donor (HD), enabling functional assessment of their tumor-killing capacity. We first evaluated different combinations of IL-15, TGF-β, and irradiated lung tumor cells (A549, E:T = 10:1), which have been reported to induce a phenotype resembling intratumoral NK cells (*18, 80*). Indeed, exposure to IL-15 together with TGF-β, A549, or a combination of both, resulted in concomitant up-regulation of CD49a and CD103 (fig. S8, A and B). The combination of IL-15 and TGF-β drove selective induction of CD39 within the CD103^+^CD49a^+^ fraction, mirroring the co-expression patterns observed in primary taNK cells (fig. S8, A and B, Fig. 3, C and D). Because the addition of A549 most closely approximates relevant aspects of the TME, we selected the three-component cocktail (IL-15, TGF-β and A549) to robustly induce the taNK phenotype over a 6-day culture period (Fig. 4D). Phenotypic validation on day 6 by flow cytometry and bulk RNA-sequencing confirmed that CD103^+^CD49a^+^ cells exhibited characteristic marker expression and a transcriptomic profile significantly enriched for the taNK signature (fig. S8, C to F) (data file S3). These in vitro-generated CD103^+^CD49a^+^ NK cells were subsequently used in a tumor cell killing assay. Sorted CD103^+^CD49a^+^ NK cells and CD103^-^CD49a^-^ NK cells populations were co-cultured with CFSE^+^ A549-b2m^ko^ target cells for 24 hours (Fig. 4D) and cytotoxicity was quantified by flow cytometry. CD103^+^CD49a^+^ NK cells displayed significantly enhanced killing capacity, confirming that this taNK-like population constitutes a dominant effector subset with potent tumor-directed cytotoxicity (Fig. 4E).

Having established the enhanced cytotoxic capacity in vitro and increased responsiveness of taNK cells ex vivo, we next asked whether this functional advantage is reflected in clinical outcomes. To evaluate the clinical relevance of taNK cells, we analyzed the association between the taNK signature and patient survival in TCGA cohorts. In LUAD, both overall NK cell infiltration and high taNK signature expression were associated with significantly improved overall survival (taNKs: HR = 0.237, *P* = 0.0022) (Fig. 4F and fig. S9A), whereas no such benefit was observed in LUSC (taNKs: HR = 1.03, *P* = 0.9) (fig. S9, A and B). Furthermore, in an independent NSCLC cohort treated with anti-PD1 (*81*), the taNK signature was significantly enriched in patients with durable clinical benefit (DCB) (Fig. 4G), suggesting a potential contribution of taNK cells in ICB responsiveness. To evaluate the broader relevance of these findings, we extended the survival analyses across additional solid tumor types. In breast cancer (BC), NK cell infiltration correlated with improved OS, while the taNK signature showed a suggestive trend (HR = 0.75, *P* = 0.06) (fig. S9A). Moreover, the taNK signature was associated with therapeutic response and T cell clonal expansion in both CRC and BC ICB cohorts, respectively (fig. S9, C to F).

Given that taNK signatures were identified as potent effector population and their signature correlate with improved survival in LUAD, we next explored their suitability for NK cell-based therapies. As a prerequisite, we explored whether taNK cells retain the ability to expand and maintain their cytotoxicity. Thus, we tested their expansion potential ex vivo using dissociated NSCLC tumor samples cultured with IL-15, K562 tumor cells or a combination of both (fig. S10A, table S2) (*82, 83*). Although combined stimulation induced the most robust expansion (fig. S10B), taNK cells expanded with IL-15 alone consistently retained the strongest cytotoxic activity against A549-b2m^ko^ cells (fig. S10C). This killing capacity was comparable to our in vitro induced taNK model (fig. S10D), demonstrating that the taNK phenotype is not a state of terminal exhaustion, but rather a functionally flexible state that can be scaled ex vivo.

Overall, CD103^+^CD49a^+^ taNK cells represent an effector subset in NSCLC with heightened cytokine sensitivity and tumor-killing capacity. Their presence correlates with improved survival in LUAD, and their ease of ex vivo expansion makes them strong candidates for future NK cell-based immunotherapies.

### ISGs and inflammation-associated regulons drive differentiation towards NK3_ENTPD1 cells

Next, we aimed to elucidate the intratumoral differentiation trajectories of taNK cells. Based on our identification of NK3_CRTAM and NK3_ENTPD1 as tumor-associated subsets, we focused our analysis on the NK3 compartment (fig. S11A). To infer their differentiation dynamics, we applied MultiVelocity (*84*), a multiomic extension of RNA velocity that incorporates chromatin accessibility to predict cell state transitions. This analysis revealed a bifurcated trajectory originating from NK3_GZMK, diverging towards NK3_CRTAM and NK3_ENTPD1 (fig. S11B). Trajectory validation using diffusion maps (Destiny (*85*)) confirmed this bifurcation and demonstrated continuous pseudotime progression along each branch (Fig. 5, A and B). Consistent trajectories and pseudotime patterns were also observed when using gene activity derived from ATAC data (fig. S11, C and D).

**Figure 5:**
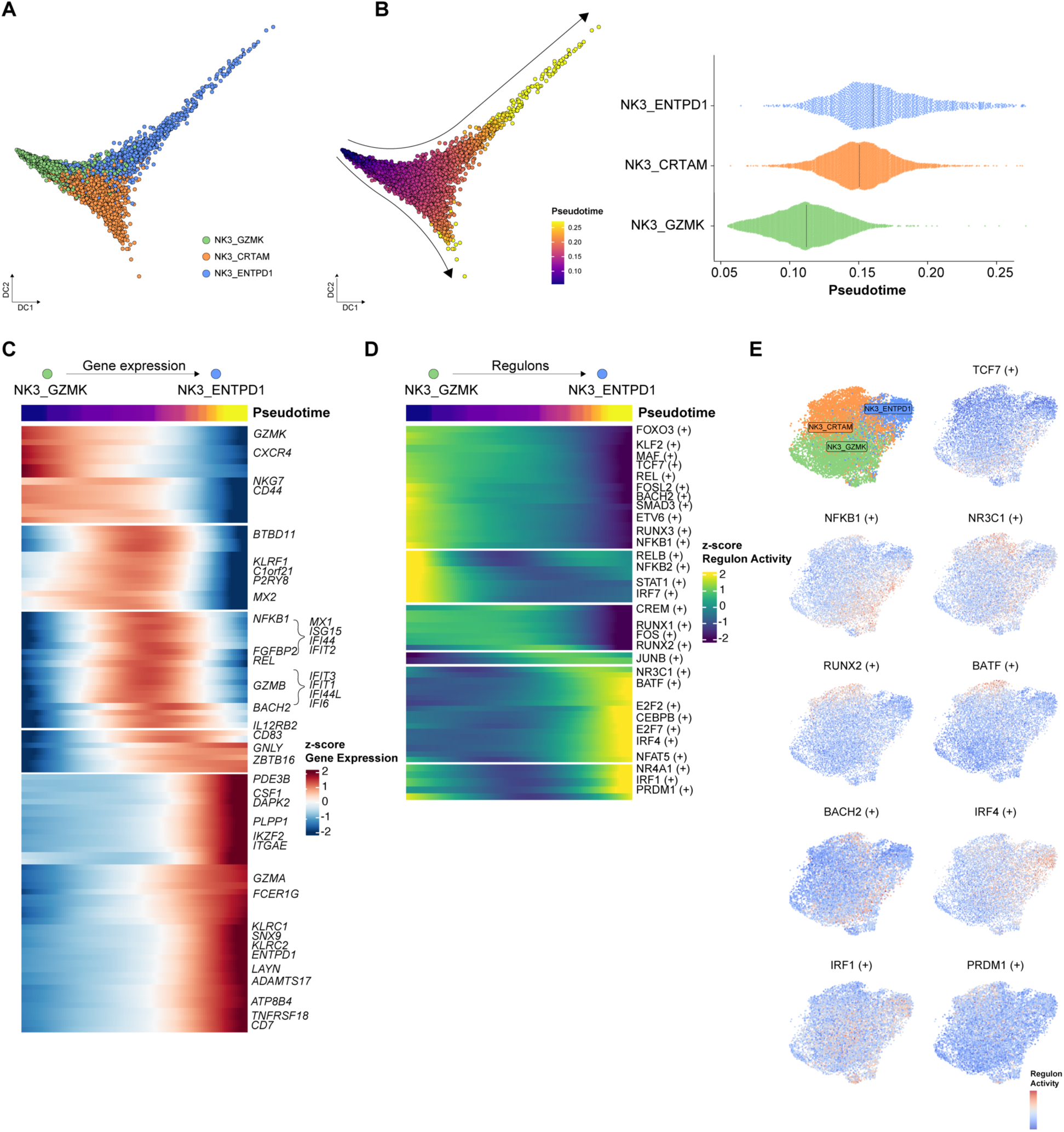
Trajectory analysis of NK3 subtypes identifies a bifurcated differentiation. **(A)** Diffusion map of single cell transcriptomes from NK3 cells, plotted along diffusion components DC1 and DC2. Cells colored by cluster (light green: NK3_GZMK; orange: NK3_CAMK4; blue: NK3_ENTPD1). (**B)** Pseudotime analysis of NK3 cells based on gene expression from RNA dataset illustrated using the diffusion-map approach (left) and group-wise stratification (right). **(C)** Dynamic heatmap showing the top 100 differentially expressed genes along the Pseudotime trajectory from NK3_GZMK to NK3_ENTPD1. Only a selection of genes displayed. **(D)** Dynamic heatmap showing enhancer-driven regulons, indicated by (+), along the Pseudotime trajectory from NK3_GZMK to NK3_ENTPD1. Only a selection of genes displayed. **(E)** UMAP displaying enhancer-driven regulons, indicated by (+), across different NK3 clusters.

Given increased expression of T_RM_-associated genes such as *ITGAE* and *ENTPD1* in NK3_ENTPD1, features linked to favorable clinical outcomes in T cells (Fig. 2, B and F), we next focused on the NK3_GZMK-to-NK3_ENTPD1 differentiation trajectory. We identified the top 100 genes exhibiting dynamic expression changes along pseudotime (Fig. 5C). Early in this trajectory, cells expressed genes associated with a CD56^bright^ phenotype including *CD44* (*11*) and *GZMK* (*12*)(*86*). This program transitioned through an interferon-stimulated phase marked by *ISGs (IFIT3, IFI44, ISG15*) and induction of *GZMB,* before culminating in the expression of genes linked to dysfunction (*KLRC1* (*87*)*, LAYN* (*88*)*, ENTDP1* (*89*)*, SNX9* (*90*)), tissue residency (*ITGAE*), adaptive NK cells (*KLRC2* (*13*)), and cytotoxicity (*GZMA*). To define the GRNs driving this progression, we performed pseudotime-based regulon analysis along the NK3_GZMK to NK3_ENTPD1 axis. This revealed a clear shift from immature transcriptional states (regulated by TCF7 and BACH2) toward inflammatory and tissue-adapted programs, including key inflammation-associated regulons (STAT1 (*91*), NFKB family (*92*)), AP-1 TFs (FOS, JUNB), and tissue-residency programs (RUNX2 (*43*)) (Fig. 5, D and E).

Altogether, integrative trajectory and regulon analyses reveal that NK3_GZMK represents an early state, giving rise to either NK3_CRTAM or NK3_ENTPD1 taNKs. The transition toward NK3_ENTPD1 is characterized by ISG activation and inflammation with the sequential engagement of adaptive, and dysfunction-associated regulatory modules, suggesting an adaptation shaped by TME-mediated cues.

### CD39^+^ tumor-associated NK cells display enhanced cytotoxicity

Because *ENTPD1* (CD39) defines a specific NK3_ENTPD1 subcluster marked by progressive acquisition of cytotoxic and inhibitory markers, we investigated whether CD39 delineates a functionally distinct taNK effector state. We first sought to provide high-resolution in situ confirmation of this population by utilizing spatial transcriptomics to visualize the co-expression of *ITGAE*, *ITGA1*, and *ENTPD1* (Fig. 6A). These three transcripts frequently co-localized within individual taNK cells, confirming the existence of this triple-positive subset within the lung TME. Quantification of *ENTPD1*^+^ taNK cells in LUAD patient LuT-722 suggests their enrichment in the tumor core and invasive margin, while absent in the adjacent non-tumor lung (fig. S12A). To validate these cells as surface-defined counterparts of NK3_ENTPD1, we next performed flow cytometric profiling. CD39^+^ taNK cells expressed higher levels of granzyme A compared to CD39^-^cells (Fig. 6B and fig. S12B). These findings suggest that CD39 marks a subpopulation with enhanced cytotoxicity within the taNK compartment.

**Fig. 6:**
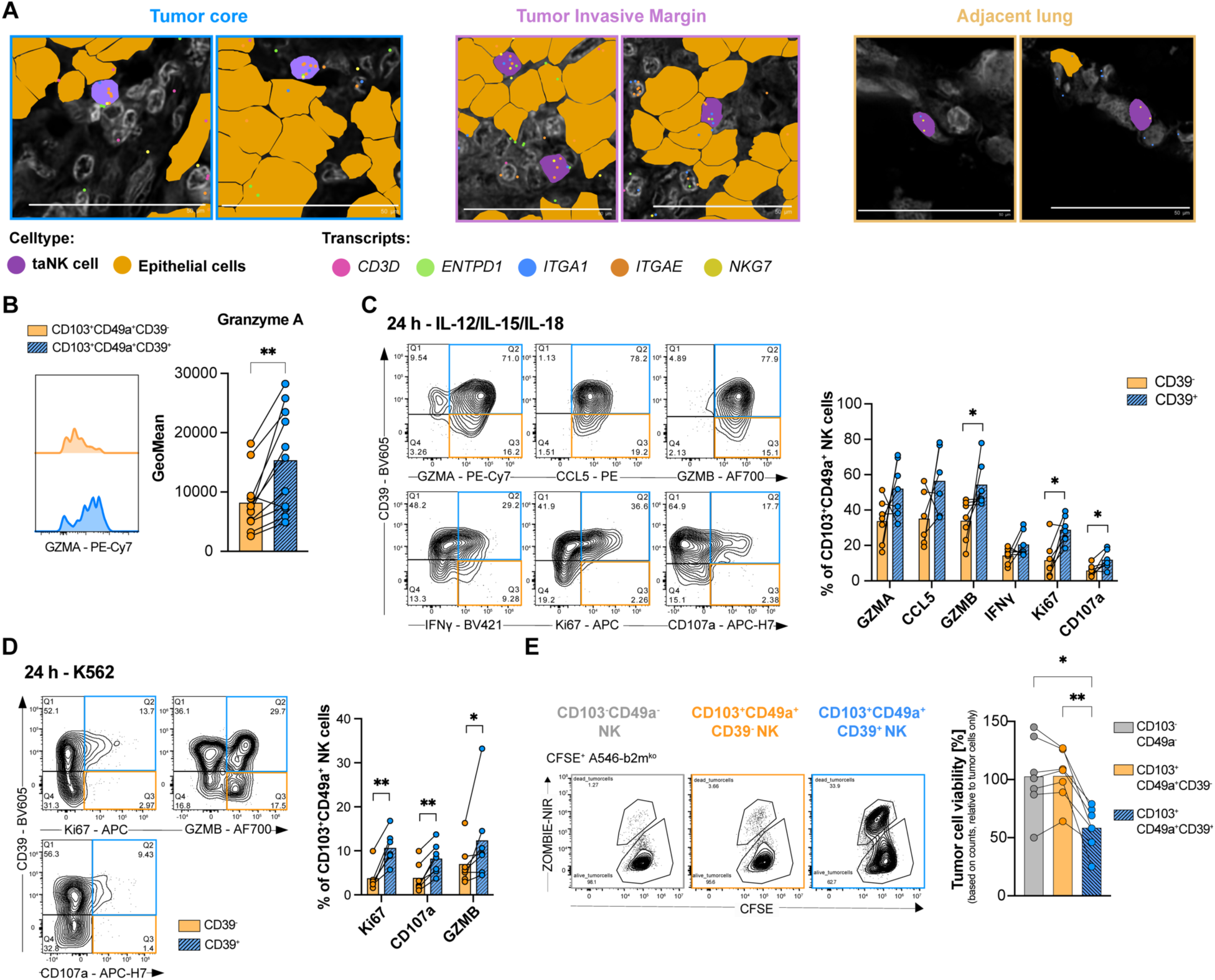
CD39^+^ tumor-associated NK cells display enhanced cytotoxicity. **(A)** Representative images from 10X Xenium experiment comparing tumor margin, core, and non-tumor areas highlighting taNK cells expressing *ENTPD1* (green), *ITGAE* (orange), *ITGA1* (blue), *NKG7* (yellow)*. CD3D* transcripts (pink) are also displayed. Scale bar: 50 µm **(B)** Representative histograms and barplot depicting the geometric mean of fluorescence intensity (GeoMean) of GZMA expression comparing CD103^+^CD49a^+^CD39^+^ (blue) with CD103^+^CD49^+^CD39^-^ (orange) NK cells from non-small cell lung cancer (NSCLC) lesions (n = 11). **(C** and **D)** Representative contour plots and quantification of indicated effector and proliferation markers in CD39^+^ and CD39^−^ subsets of CD103^+^CD49^+^ NK cells from non-small cell lung cancer (NSCLC) lesions following 25 hours stimulation with (C) IL-12/IL-15/IL-18 or (D) K562 for 24 hours (n = 7). **(E)** Representative plots of ZombieNIR^+^CFSE^+^ A549-b2m^ko^ tumor cells after 24 hours co-culture with sorted CD103^-^CD49a^-^, CD103^+^CD49a^+^CD39^-^, or CD103^+^CD49a^+^CD39^+^ NK cell from the in vitro taNK cell platform (left). Tumor cell viability was determined by absolute counts, normalized to a tumor-only control and represented as bar graphs (mean with individual samples) (n = 7). Data were analyzed by (B, C and D) paired Wilcoxon signed-rank test and (E) non-parametric paired Friedman test with Dunn’s multiple comparisons. *\*P < 0.05, **P < 0.01, ***P < 0.001, ****P < 0.0001*

We then assessed functional responsiveness by stimulating taNK cells from NSCLC tumor lesions for 5 and 24 hours with either IL-12/IL-15/IL-18 or co-cultured them with K562 tumor cells (Fig. 4A and 6, C and D, fig. S12, C and E, table S2). After 5 hours, CD39⁺ taNK cells showed a significantly higher frequency of Ki67⁺ cells under both conditions, indicating a greater proliferative potential associated with CD39 expression (fig. S12, C and D). After 24 hours, CD39^+^ taNK cell population exhibited enhanced expression of granzyme B, Ki67, and CD107a, highlighting superior functional capacity (Fig. 6, C and D). Under cytokine stimulation, we also observed a patient-dependent trend toward higher expression of CCL5, granzyme A, and IFNγ in CD39^+^ taNK cells, a pattern less evident under K562 co-culture conditions (Fig. 6, C and D, fig. S12E). To determine whether these functional differences translate into enhanced tumor killing, we utilized our in vitro taNK model to directly compare the cytotoxic capacity of CD39^+^ CD103^+^CD49a^+^ NKs, CD39^-^ CD103^+^CD49a^+^ NKs, and CD103^-^CD49a^-^ NK cells (Fig. 4D). Sorted populations were co-cultured with labeled A549-b2m^ko^ target cells and cytotoxicity was quantified by flow cytometry. CD39^+^ CD103^+^CD49a^+^ NKs exhibited a significantly higher killing capacity compared to the other subsets, establishing them as the dominant cytotoxic effector subset with within the taNK compartment (Fig. 6E).

Thus, CD39⁺ taNK cells represent a functionally distinct and dominant cytotoxic effector population within the NSCLC TME.

### NKG2A blockade enhances the effector capacity of CD39^+^ taNK cells

Since *KLRC1* (NKG2A) expression is significantly enriched in taNK cells (Fig. 2, B and F), we hypothesized that this inhibitory axis may restrain their full effector capacity. Co-expression analysis of our single nucleus data confirmed that *KLRC1* transcripts were most strongly enriched with *ENTPD1* (CD39), *ITGAE* (CD103), and *ITGA1* (CD49a) within the NK3_ENTPD1 cluster, with lower expression observed in NK3_CRTAM (fig. S12F). We further validated this signature at the protein level using flow cytometry, which revealed significantly higher NKG2A expression on CD39^+^ taNK cells compared to their CD39^-^ counterparts (Fig. 7A). Given this this selective enrichment, we next asked whether anti-NKG2A blockade could further potentiate the heightened cytotoxic capacity. To address this, we employed the patient-derived tumor fragment (PDTF) platform, an ex vivo 3D model that preserves the native cellular architecture and immune microenvironment of human lung tumors (*93*). PDTFs from NSCLC patients (n = 7, table S2) were treated for 48 hours with isotype control or anti-NKG2A, a biosimilar to Monalizumab, a monoclonal antibody under investigation in phase 3 trials for NSCLC (NCT05221840). Post-treatment, PDTFs were pooled by condition, digested, and analyzed by flow cytometry (Fig. 7B). Anti-NKG2A treatment resulted in a reduction of NKG2A surface expression on both, CD39^+^ and CD39^-^ taNK cells, indicating effective target engagement (fig. S12G). Furthermore, anti-NKG2A treatment led to a significant increase in the proportion of CD39⁺ NK cells expressing GZMB or GZMA, while the CD39⁻ subset remained unchanged (Fig. 7C, fig. S12, H and I). When specifically gating on CD39⁺ cells, only GZMA showed a significant increase, suggesting a preferential up-regulation of GZMA upon anti-NKG2A (fig. S12J). These findings identify CD39⁺ taNK cells as a tumor-enriched NK cell population that is responsive to NKG2A blockade. Anti-NKG2A therapy either expands existing CD39⁺GZMA⁺ and CD39⁺GZMB⁺ populations or promotes their induction from CD39⁻ precursors, thus enhancing their cytotoxic potential.

**Fig. 7:**
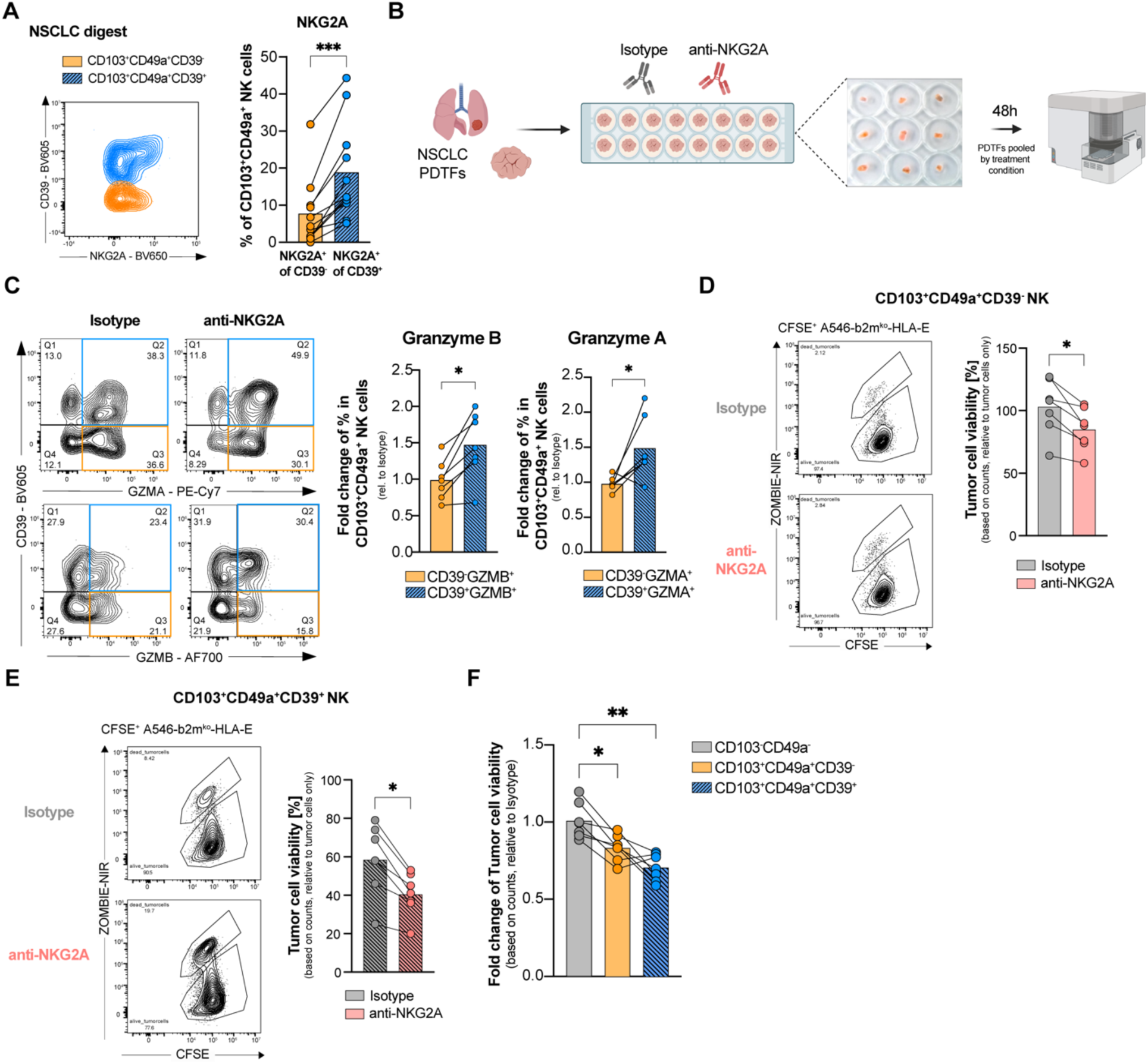
NKG2A blockade enhances the effector profile and cytotoxicity of CD39^+^ tumor-associated NK cells. **(A)** Representative flow cytometry plot and quantification of NKG2A expression on CD39^+^ versus CD39^−^ subsets of CD103^+^CD49a^+^ NK cells from non-small cell lung cancer (NSCLC) digests (n=11) . **(B)** Experimental workflow for the Patient-Derived Tumor Fragment (PDTF) assay and anti-NKG2A treatment. (Graphics created with Biorender.com (*145*)) **(C)** Representative contour plots and bar plots showing percentage of GZMB^+^ and GZMA^+^ populations co-expressing CD39 (dashed blue) and their CD39^-^counterparts (orange) in CD103^+^CD49a^+^ NK cells from PDTFs upon anti-NKG2A treatment for 48 hours illustrated as fold change to isotype control (n=7). (**D** and **E)** Representative flow cytometry plots and corresponding quantification of tumor cell viability (A549-b2mko-HLA-E) following co-culture with sorted (D) CD39^−^ or (E) CD39^+^ in vitro induced taNK cells treated with isotype or anti-NKG2A. Tumor cell viability is illustrated as a percentage relative to the tumor cells only control (n = 6). **(F)** Mean fold change of tumor cell viability in anti-NKG2A-treated conditions relative to their respective isotype controls across indicated *in vitro* induced NK cell subsets (n = 6). Data were analyzed by [(A), (C), (D) and (E)] paired Wilcoxon signed-rank test, (F) non-parametric paired Friedman test with Dunn’s multiple comparisons. *\*P < 0.05, **P < 0.01, ***P < 0.001, ****P < 0.0001*

To determine whether the increased granzyme expression translates into enhanced tumor killing, we utilized our in vitro taNK model, which similarly displays the highest NKG2A surface levels within the CD39^+^ CD103^+^CD49a^+^ NK population (fig. S12K). Sorted CD39^+^ or CD39^-^CD103^+^CD49a^+^ NKs were co-cultured with A549-b2m^ko^-HLA-E target cells in the presence of either an isotype control or anti-NKG2A antibody (fig. S12L).

NKG2A blockade significantly enhanced the killing capacity of both CD39^+^ and CD39^-^CD103^+^CD49a^+^ NK subsets compared to isotype controls, whereas tumor cell viability remained unchanged in co-culture with CD103^-^CD49a^-^ NK cells (Fig. 7, D and E, fig. S12M). When tumor cell viability was normalized as fold-change relative to the isotype control, the CD39^+^ subset emerged as the most potent effector population (Fig. 7F). Although the difference in killing between CD39^+^ and CD39^-^ CD103^+^CD49a^+^ NKs did not reach statistical significance, the CD39^+^ subset consistently mediated the strongest reduction in tumor cell viability (*P* < 0.01 vs. CD103^-^CD49a^-^).

Collectively, these results indicate that CD39^+^ taNKs represent a highly responsive effector state that achieves elevated tumor-directed cytotoxicity upon NKG2A blockade.

## DISCUSSION

Despite increasing efforts to harness NK cells for cancer immunotherapy (*9*), the transcriptional and functional states of intratumoral NK cells remain incompletely defined. Here, we provide an integrated snRNA-seq and snATAC-seq atlas of NK cell states in human lung cancer. We identify two taNK clusters co-expressing *ITGAE* (CD103) and *ITGA1* (CD49a) that display markers of dysfunction (*ENTPD1, TOX, KLRC1*) yet retain a strong effector program (*CCL5, GZMA*) and enhanced cytotoxic capacity. In particular, we defined a CD39⁺ subpopulation of (CD103⁺CD49a⁺) taNKs that likely emerges from *GZMK*⁺ NK cells in response to inflammatory signals. Crucially, our functional assays identify this CD39⁺ subset as the predominant NK effector population within NSCLC, exhibiting superior killing capacity. Furthermore, this population is the primary responder to anti-NKG2A treatment, which selectively potentiates their cytotoxicity. Together, these findings establish CD39⁺ taNKs as critical anti-tumor effectors and a targetable NK cell state in NSCLC.

Although NK cell dysfunction within tumors is well-established (*17*), the terminology of ’tumor-associated NK cells’ (TaNKs) was only recently introduced by Tang *et al.* (2023) to describe a cluster characterized by a stress-related transcriptional profile identified through pan-cancer integration correlating with poor prognosis (*26*). In contrast, the CD103^+^CD49a^+^ taNKs we identify here represent a distinct lineage characterized by a tissue-residency program showing increased anti-tumor cytotoxicity. While our taNKs share certain markers associated with dysfunction, the coordinated co-expression of tissue-residency markers (CD103, CD49a, CXCR6) has only recently been recognized across cancer types (*21, 23, 25*). Our findings indicate that despite this “dysfunctional” profile, human taNKs retain substantial functional plasticity. Utilizing an in vitro model incorporating IL-15, TGF-β, and tumor cell exposure, factors known to drive tissue-resident differentiation (*18, 80*), we recapitulated the taNK phenotype and confirmed their enhanced cytotoxic capacity. This suggests that taNK cells do not represent a terminally exhausted population, but rather a specialized effector differentiation stage. This interpretation is consistent with murine studies showing that NK cells expressing inhibitory receptors can regain function following stimulation (*94*). Nonetheless, future work will be needed to determine whether this state remains or becomes fixed under chronic tumor-imposed pressure.

The clinical relevance of taNK cells is supported by the association of the taNK signature with improved survival in LUAD, a benefit not observed in LUSC. While the mechanism remains unclear, this divergence may reflect context-specific features of the TME. For example, the taNK signature may be functionally constrained by a more stromal- and myeloid-rich LUSC microenvironment (*95, 96*). Compared with LUAD, LUSC has been associated with increased extracellular-matrix remodeling and suppressive immune populations, which may limit NK cell effector function despite detectable transcriptional signatures (*95–97*). However, this interpretation remains speculative and will require future spatial studies. Furthermore, the ability to expand taNK cells ex vivo while preserving their potent cytotoxicity further highlights their promise for adoptive NK cell-based therapies. Together, our results reinforce the concept that NK cell dysfunction is not irreversible but dynamically shaped by the tissue context, and they position taNK cells as clinically relevant populations in lung cancer.

Trajectory analysis revealed a bifurcated differentiation route from *GZMK*^+^ NK cells progressing toward two tumor-associated states: an *ENTPD1*^+^ and a *CRTAM*^+^ state. While their precise cellular origin remains unresolved, both circulating and pre-existing tissue-resident NK cells may contribute. Recent murine studies suggest that resident-memory NK cells can persist in tissues long after infection clearance (*98–101*). In the human lung, Brownlie *et al.* described a hyperresponsive, adaptive-like NK cell population marked by elevated CD49a expression (*79*). As *CRTAM*⁺ NK cells exhibit stronger enrichment for adaptive NK cell signatures and only partially enrich in tumor tissue (Fig. 2E and fig. S3H), this subset may reflect a tissue-resident, adaptive-like subset that expands upon tumor progression. This interpretation is supported by our cross-modality analysis, which suggests distinct epigenetic origins for the *ENTPD1*⁺ and *CRTAM*⁺ lineages (fig. S3, D to G). However, this interpretation remains speculative and warrants further investigation. In contrast, *ENTPD1*⁺ NK cells predominate in tumors and its transition toward a CD39⁺ state is marked by ISG activation, cytotoxicity, and inflammatory regulators. This progression involves acquisition of tissue-residency features and is further shaped by up-regulation of GRNs linked to dysfunction (*NR4A*) (*73*), proliferation (E2Fs), cytotoxicity, and adaptive-like immunity (*IRF4, PRDM1*). Of note, many of these regulons are shared with exhausted or tumor-reactive T cells (*102, 103*). For instance, NR4A1 drives T cell exhaustion via suppression of AP1-mediated activation (*74*), while IRF4 is implicated in both, NK and T cell activation and effector function (*75, 76, 104*). These parallels suggest that NK and T cells may converge on shared transcriptional programs in response to an inflammatory tumor milieu.

Functionally, this CD39⁺ taNK subset exhibits elevated levels of granzyme A, NKG2A, and proliferation markers, displays potent anti-tumor cytotoxicity and shows a heightened effector profile following cytokine stimulation. In accordance with our findings, granzyme A and CD39 have been reported to be enriched on NK cells within the tumor core of NSCLC, where they also show increased Ki-67 induction upon restimulation (*79*). Thus, although CD39 is an ectoenzyme within the adenosine pathway potentially involved in local immunosuppression, our data support its use as a marker of a taNK effector subset (*105*). These cells may engage immune checkpoint pathways as an adaptive feedback mechanism, potentially limiting excessive effector activity within the chronically stimulatory TME (*106, 107*). Their resemblance to CD103^+^CD39^+^ tumor-reactive CD8 T_RM_ cells, which are functionally responsive despite immune checkpoint expression and are enriched following ICB (*67–69, 108–111*), raises the question of how NK cells acquire transcriptional features typically attributed to TCR-dependent signaling. Defining additional receptors-ligand interactions or cytokine cues that shape these shared effector states will be an important direction for future work.

Therapeutically, our PDTF and in vitro models demonstrate that NKG2A blockade enriches the CD39⁺ taNK compartment, likely through expansion of pre-existing effectors and potentially inducing their differentiation from CD39⁻ taNK cells. Although CD39^+^ taNK cells already exhibit strong baseline cytotoxicity, disrupting the NKG2A-HLA-E axis further amplifies their tumor killing capacity, identifying this subset as a responsive therapeutic target. Beyond NKG2A, their co-expression of multiple inhibitory receptors suggests additional checkpoint blockade strategies targeting TIGIT, LAG3, or CD39 itself. Indeed, CD39 inhibition has been shown to enhance intratumoral NK cell infiltration (*112*) and promote T cell activation and proliferation (*113*). These findings are supported by clinical trials using anti-CD39 showing encouraging activity, particularly in combination with PD-1 blockade (*114, 115*). Incorporating NK-focused immune monitoring into these studies may provide important insights into taNK modulation. Furthermore, taNKs express activating receptors such as NKG2D (*KLRK1*) and CD28H (*TMIGD2*), positioning them as attractive candidates for multi-specific NK cell engagers (NKCEs) (*9*). We propose that NKCEs incorporating an IL-15 linker may be particularly effective, as our data show that taNK cells can be expanded using IL-15 while maintaining potent cytotoxic function, supporting improved proliferation, and persistence of this effector pool.

While PDTFs preserve critical aspects of tumor architecture and immune interactions, they remain a semi-physiological ex vivo system that cannot fully capture systemic influences (*93*). Nevertheless, the consistent identification of CD39⁺ taNK cells as dominant effectors across our experimental models provides a robust functional framework. Future in vivo studies will be essential to validate these dynamics in a complete physiological setting and to assess the durability of taNK cytotoxicity following therapeutic intervention.

In conclusion, we here provide an integrated transcriptional and epigenetic atlas of NK cell states in human lung cancer and identify CD39^+^ tumor-associated NK cells as a highly cytotoxic effector subset. We further show that their anti-tumor activity can be selectively potentiated by NKG2A blockade, establishing a functional link between taNK phenotypes and therapeutic responsiveness. Together, these insights highlight the CD39^+^ taNK axis as a promising target for next-generation NK-based immunotherapy in lung cancer.

## MATERIALS AND METHODS

### Study design

This study aimed to investigate the functional specialization and heterogeneity of intratumoral natural killer (NK) cells in treatment-naïve patients with non-small cell lung cancer (NSCLC) using a multiomic single-nucleus approach (snRNA-seq and snATAC-seq) alongside detailed functional assays. Patients with either lung adenocarcinoma (LUAD) or lung squamous cell carcinoma (LUSC) across different stages, including both sexes and smokers and non-smokers, were selected, and samples were prescreened to ensure sufficient NK cell numbers for single-nucleus profiling. Key findings from single-nucleus profiling were further characterized at the protein level and functionally validated using 2D assays, an in vitro tumor-associated NK (taNK)-mimetic platform, and 3D patient-derived tumor fragment (PDTF) co-cultures. Across these platforms, we employed multiparameter flow cytometry to establish baseline phenotypes and evaluate therapeutic relevance through restimulation and cytotoxicity read-outs in both NSCLC and non-tumor lung samples. Investigators were not blinded to group assignments during experiments or analyses. Although statistical methods were not used to pre-determine sample size, sample sizes were chosen based on previously published results. The number of biological replicates varied by experiment based on sample availability and is explicitly stated in the figure legends. Detailed information of NSCLC patients used for the study can be found in table S1 and table S2. To ensure data robustness and minimize technical noise, sub-gating was only performed on samples where populations were sufficiently well-defined to allow for consistent and reproducible gating.

### Tumor sample collection

Fresh tumor samples were collected from NSCLC patients undergoing primary surgical treatment at University Hospital Basel or Kantonsspital Baselland Liestal, Switzerland. The study was approved by the local Ethical Review Board (Ethikkommission Nordwestschweiz, EK321/10), performed in compliance with all relevant ethical regulations, and all patients consented in writing to the analysis of their tumor samples. Patient characteristics are summarized in table S1 and S6.

### In vitro taNK cell platform

Primary human NK cells were negatively isolated from healthy donors PBMCs using EasySep Human NK Cell Isolation Kit (StemCell, #17955). After isolation, NK cells were cocultured with irradiated A549 (RRID: CVCL_0023; 30 Gy) at a ratio of 10:1 in NK cell culture medium (RPMI 1640 (Brunschwig, #CPCRPMI-A) supplemented with 1 mM of sodium pyruvate (Sigma-Aldrich, #11360070), 1X Minimum Essential Medium (MEM) nonessential amino acids (Gibco, #11140050), 2 mM HEPES (Gibco, #15630080), 10% heat-inactivated human serum (Blood Bank, University Hospital Basel, Switzerland), 1% penicillin–streptomycin (Gibco, #15140122) and 50 µM β-mercaptoethanol (Gibco, # 11528926)) supplemented with 10 ng/ml IL-15 and 10 ng/ml TGF-ß. On 3 days, medium was refreshed and fresh irradiated A549 (RRID: CVCL_0023; 30 Gy) were added. After 6 days of culture, cells were sorted (ZombieNIR^−^, CD56^+^CD3^-^, either CD103^+^CD49a^+^, CD103^+^CD49a^+^CD39^+^, CD103^+^CD49a^+^CD39^-^ or, CD103^-^CD49a^-^) using Cytek Aurora Cell Sorter.

### Functional assays

For cytokine restimulation assays, whole tumor lesions were incubated for 5 or 24 hours either alone, at a ratio of 1:1 with the irradiated target cell line K562 (RRID: CVCL_0004; 90 Gy), or with a cytokine cocktail (IL-12: 1 ng/ml, PeproTech, #200-12; IL-15: 10 ng/ml, PeproTech, #200-15; IL-18: 5 ng/ml, Gibco, #PHC0186) at a density of 1x10^6^ cells/ml per well in NK cell culture media. Four hours before harvest, a media mix consistent of CD107a-APC-H7 (1:150) (Biolegend, H4A3, #328630), Brefeldin A (1:1000) (eBiosciences, #00-4506-51), and Monensin (1:1000) (eBiosciences, #00-4505-51) was added to block cytokine secretion. Subsequently, cells were harvested and stained for flow cytometry. For killing assays, cells were co-cultured at a ratio of 2:1 with CFSE-labelled A549-b2m^ko^ (RRID: CVCL_C3CC) or A549-b2m^ko^-HLA-E (RRID: CVCL_C3CC) at a density of 1x10^6^ cells/ml per well in NK cell culture medium. After 24 hours of incubation, counting beads were added and alive tumor cells were counted using flow cytometry. In some assays, anti-NKG2A (10 µg/ml, Ichorbio, ICH5017), or Isotype control (human IgG4, QA16A15: 10 µg/ml, Biolegend, #403701) were added during incubation.

### Fluorescence-activated cell sorting for the single-nucleus experiment

Tumor lesions were thawed, washed with PBS, and stained according to the flow cytometry protocol mentioned above. For live/dead staining the Zombie NIR fixable viability kit (Biolegend, 1:1000) was used. After human Fc receptor binding inhibitor (Invitrogen) incubation, cells were stained in staining buffer (PBS + 1% bovine serum albumin (BSA) (Sigma; #10735086001)) containing antibodies (CD45-BV711: H130, Biolegend (#304050), 1:100; CD8-BV421:SK7, Biolegend (#344748), 1:100; CD8-APC: SK1, Biolegend (#344722), 1:100; CD19-FITC: HIB19, Biolegend (#302206), 1:100; CD14-FITC: M5E2, Biolegend (#982502), 1:100; CD56-PE: 5.1H11, Biolegend (#981202), 1:100). Cells were washed twice with staining buffer. Subsequently, Fluorescence-activated cell sorting (FACS) was performed using BD Melody or BD FACS Aria III. Cells were gated on single cells, live (Zombie NIR^-^), CD45^+^, CD19^-^CD14^-^ and CD56^+^CD3^+^. Cells were sorted in cold PBS with 2% FBS (Pan Biotech) and used immediately for the single cell experiment.

### Single cell library preparation and sequencing

Sorted single cell suspensions were further processed using the Chromium Next GEM Single Cell Multiome ATAC + Gene Expression protocol (CG000338, Rev C) provided by 10x Genomics according to the manufacturer’s instructions. First, nuclei Isolation was performed using the Nuclei Isolation protocol for primary cells for Single Cell Multiome ATAC + Gene Expression Sequencing (CG000365, Rev B) provided by 10x Genomics. Most reagents were used as recommended by indicated protocol (BSA (Sigma, #10735086001), Nonidet P40 substitute (Sigma, #98379), Tween-20 (Sigma, #93773), DTT (Sigma, #43816)). Lysis was performed using the lysis buffer at 0.5X for 3 min. Subsequently, nuclei transposition was conducted and nuclei were loaded in a 10x Genomics Chip J to achieve a target recovery of approximately 10’000 cells for each sample. Cell-barcoded 3′ gene expression (GEX) libraries and ATAC libraries were sequenced using the Illumina NovaSeq6000 system at recommended sequencing depths (GEX: 20’000 reads per nucleus, ATAC: 25’000 reads per nucleus). GEX and ATAC libraries were aligned and annotated with CellRanger ARC v.2.0.2 using the human reference genome GRCh38 2020-A-2.0.0 provided by 10x Genomics.

### Single cell data analysis

The read count matrices were processed and analyzed using R v4.3.0 or higher, Signac v1.14.0 (*28*), and Seurat v5.1.0 (*27*), with provided standard workflows using default parameters for all functions unless otherwise specified. Peak calling was performed using MACS2 v2.2.7.1 (*116*). Peak sets called from different donors were reduced to a joint peak set. For quality control, cells were filtered based on ATAC and GEX parameters (ATAC: nFeatures > 200, nCount > 1’000 & < 100’000, nucleosome signal < 2, TSS enrichment > 1, blacklist fraction < 0.05, fraction of reads in peaks per cell (FRiP) > 0.25; GEX: nFeatures > 200, nCount > 1’000 & < 25’000, percent mitochondrial genes < 20, percent ribosomal genes > 0.05). Next, GEX count matrices were log-normalized, scaled and integrated using canonical correlation analysis (CCA) in order to remove donor-specific alterations. Further, peak matrices were normalized, and donor-specific batch effects were corrected by projecting into a low dimensional space (LSI). Subsequently, WNN integration was performed in order to integrated RNA and ATAC modalities using the integrated dimensional reductions. After clustering and UMAP representation, differential gene expression (DEG) analyses were conducted with the Seurat implemented function *FindAllMarkers* using the MAST (*117*) test. Clusters corresponding to regulatory T cells (*FOXP3*, *CD4*, *CTLA4*, *IL2RA*), B cells/plasma cells (*CD79A, MZB1, JCHAIN*) and monocytes (*CST3*, *FCER1A*, *XCR1*, *LYZ, CD86*) were removed to clean the dataset. Moreover, a signature score for T cells (*CD3D*, *CD3G*) was calculated using the *AddModuleScore_UCell* function from the *UCell* package (*118*) (v2.6.2) in order to identify T cells. Cells with a signature score higher than 0.05 were removed from the dataset in order to obtain a clean NK cell dataset. Subsequently, aforementioned steps (normalization, scaling, integration, dimensional reduction, and clustering) were repeated for the purified NK cell dataset. For scaling, percentage of mitochondrial genes were regressed. Additionally, genes irrelevant for clustering, such as histone genes and ribosomal genes were excluded from variable features using a gene blacklist from Zheng *et al.* (*119*). For clustering and UMAP calculations, the first 15 principal components (PCs) were used for the RNA dataset and the first 20 PCs for the ATAC dataset based on variance calculations. Cluster annotations and subset identifications were performed using DEG analysis and calculations of signature scores from publicly available datasets (“dysfunctional” (*66*), “exhausted” (*65*), “cytotoxic” (*65*), “tumor-reactive” (*54*), “resident-memory” (*119*), “terminally exhausted” (*119*) CD8, ILC signatures (*30*), NK signatures(*11, 13, 31*), stress signature (*26*)) using the *UCell* package (v2.6.2) and the escape v1.12.0 (*120*) package. Gene counts were aggregated using Seurat-implemented function *AggregateExpression.* For the heatmap visualization of gene transcripts of aggregated counts, only transcripts were selected that were present in more than 10% of the cells of at least one cluster. *ClusterProfiler* (v4.4.2) (*121*) was used to perform gene set enrichment analysis (GSEA). The aforementioned analysis steps were repeated on the subset of NK3 cells, beginning with dataset normalization.

### Regulon analysis (SCENIC+)

Briefly, SCENIC+ (*33*) is a recent development of the SCENIC tool that takes advantage of multiomic data. It predicts genomic enhancers along with candidate upstream TF and links these enhancers to candidate target genes. Specific TFs for each cell subset are thus predicted based on the concordance of TF binding site accessibility, TF expression, and target gene expression as contained in multiomic data (scRNA-seq and scATAC-seq).

For SCENIC+ analysis, scRNA-seq data was preprocessed using Seurat as previously described and scATAC-seq data was processed using pycisTopic (v2.0a0) (*33*). We filtered cells based on both, scRNA-seq (as described above) and scATAC-seq analysis (thresholds were automatically defined by the *pycistopic qc* function). A model of 40 topics was selected for both, the complete NK cells dataset and the NK3 cells only. PycisTopic (*33*) (v2.0a0) was used to binarize the topics and identify differentially accessible regions between cell types. A custom cisTarget database was then generated based on the consensus peak detected in our datasets. Next, *pycistarget* (*33*)was applied to identify enriched motifs in the identified candidate enhancer regions using the custom cisTarget database. Finally, enhancer-driven gene regulatory networks were inferred using SCENIC+ (v1.0a1) (*33*). Regulon specificity scores (RSS) were computed using the Jensen-Shannon Divergence (*122*). When exploring the inferred regulons within the entire NK cell dataset or the NK3 cells only, we identified 104 and 125 active TFs, among which 47 and 58 were activators (i.e., associated with increased chromatin accessibility and RNA expression of the target genes), respectively.

### MultiVelocity analysis

MultiVelo (*84*) (v0.1.5) was used to infer velocity trajectory based on both RNA expression and ATAC. Briefly, MultiVelo employs a probabilistic latent variable model to estimate gene regulatory switch times and rate parameters, offering a quantitative framework to capture the temporal dynamics between epigenomic and transcriptomic changes. When applied to single-nucleus multiomic datasets, MultiVelo reveals two distinct chromatin accessibility–driven regulatory mechanisms, quantifies concordance or discordance between transcriptomic and epigenomic states at the single-nucleus level, and infers time lags between these molecular layers.

The analysis was performed on the NK3 subset pooled across all samples. For the RNA part, low quality cells were filtered based on their RNA (minimal number of counts = 300, maximum number of counts = 4000). Data were normalized and only the top 1000 variable genes were retained for downstream analysis. For the ATAC analysis, peaks were aggregated using the *aggregate_peaks_10x* function. Outlier cells were removed based on their total number of detected peaks, with a minimum threshold of 2,000 and a maximum of 35,000. Only cells that passed both filtering steps were retained for downstream analysis. Chromatin accessibility data were smoothed using the *knn_smooth_chrom* function, and WNN neighbors computed with Seurat were used in subsequent steps. The joint chromatin-RNA model was then optimized using the *recover_dynamics_chrom* function, followed by computation of the velocity graph with *velocity_graph*, both using default parameters.

### Statistical analysis

The R package ggsignif (*123*) v0.6.4 and GraphPad Prism v10.3.0 were used to calculate statistical significances. These analyses included Wilcoxon signed-rank test, non-parametric Kruskal-Wallis and Friedman test with post-hoc Dunn test, and paired Wilcoxon signed-rank test as described in the Fig. legends. *P* values > 0.05 were considered not significant, and *P* values < 0.05 were considered significant: \**P* < 0.05, \*\**P* < 0.01, \*\*\**P* < 0.001, and \*\*\*\**P* < 0.0001.

## Supplementary Materials

Supplementary Materials and Methods

figs. S1 to S12

Tables S1 to S3

Data files S1 to S4

## Supporting information

Whole Supplementary Material

## Acknowledgements

We thank the patients for allowing us to report their clinical information and data. We thank Dr. R. Ivanek, Dr. J. Roux and Dr. I. Cervenka from the DBM Bioinformatics Core Facility for support and feedback with bioinformatics. Furthermore, we thank the Pathology Department of the University of Basel, especially Dr. A. Pant and Dr. S. Uzun. Additionally, we would like to thank the Flow Cytometry Facility of the University of Basel, and the Genomics Facility Basel of the University of Basel and the Department of Biosystems Science and Engineering, ETH Zurich for the library sequencing. Computational data processing for this project were performed at sciCORE (http://scicore.unibas.ch/) scientific computing center at the University of Basel. We used ChatGPT (GPT-4 and GPT-5 mini) and Google Gemini (2026) to assist with refining sentence structure and phrasing. All content was reviewed and verified by the authors.

## Authors contributions

Bioinformatics Investigation: C.Serger, L.R., M.T.S., W.R., and E.C. Conceptualization: C.Serger, M.P.T., M.N., K.S., A.R., E.V., and A.Zippelius. Design and Interpretation: C.Serger, I.F., L.R., M.T.S., S.T., T.T.L., M.N., K.S., E.V., and A.Zippelius. Funding acquisition: M.B., H.L., C.F., E.V., A.R., and A.Zippelius. In vitro Investigation: C.Serger, P.H., A.B., C.A.P. and N.O. Methodology: P.H, C.Schultheiß, and A.Zingg. Resources: A.H., D.L., L.N., K.M., and M.S.M. Supervision: M.P.T., M.N., C.F., K.S., E.V., and A.Zippelius. Writing-original draft: C.Serger. Writing-review and editing: C.Serger, L.R., M.T.S., I.F., A.R., K.S., E.V., and A.Zippelius.

## Data and materials availability

Gene expression matrices from single-nucleus Multiome sequencing are available in the Gene Expression Omnibus (GEO) under GSE304741. 10X Xenium Spatial Transcriptomics data are available under GSE319755. Raw sequencing FASTQ files from single-nucleus Multiome sequencing are deposited in the European Genome-Phenome Archive (EGA) under accessions EGAS50000001189 and EGAS50000001190. Raw sequencing FASTQ files from bulk RNA sequencing will be available through the EGA under EGAS50000001658 and EGAD50000002387. Due to data security guidelines of the University of Basel, raw sequencing data are under restricted access. Researchers can apply for access by contacting the Data Access Committee (DAC) at. All the custom code used in this study are provided on GitHub (*139–140*) and permanently stored at Zenodo (*141*). All materials are commercially available as described in Materials and Methods. Raw tabulated data underlying the figures are provided in data file S4.

