## Supplementary material for "CD39 defines a cytotoxic tumor-associated NK cell state responsive to NKG2A blockade in lung cancer": Whole Supplementary Material

Science AAAS

Clara Serger *et al.*

**The PDF file includes:**

Supplementary Materials and Methods
Figs. S1 to S12
Tables S1 to S3

### SUPPLEMENTARY MATERIALS AND METHODS

#### Isolation of human PBMCs and tumor-infiltrating lymphocytes

Resected tumor lesions and PDTFs were enzymatically dissociated into single cell suspensions using Accutase (Innovative Cell Technologies, #ICTAT104-100), collagenase type IV (1mg/ml, Worthington, #LS004189), hyaluronidase (1mg/ml, Sigma-Aldrich, #H6254-1G), and DNase type I (40ug/ml, Sigma-Aldrich, #D5025). Subsequently, cells were washed in 1x phosphate-buffered saline (PBS) and filtered using a 100 µm filter mesh. Human peripheral blood mononuclear cells (PBMCs) were isolated by density gradient centrifugation, using Histopaque-1077 (Sigma-Aldrich, #10771). Resulting single cell suspensions (tumor lesions) were cryopreserved in fetal bovine serum (FBS) (Brunschwig, #FBS-LE-12A) and 10% dimethyl sulfoxide (Sigma, #D2650), and stored in liquid nitrogen until further usage. The dissociation protocol was optimized to preferentially preserve immune cell populations, tumor cells are less likely to remain viable following the enzymatic digestion and subsequent cryopreservation.

#### Patient-derived tumor fragment (PDTF) assays

Obtained tumor lesions collected for PDTF cultures were cut into small tumor fragments of 1 – 2 $mm^3$ size on ice as reported in Roelofsen *et al. (124)*. Subsequently, single PDTFs were mixed to ensure uniform representation of the tumor lesions and were cryopreserved until further use. For the assays, individual PDTFs were embedded in artificial extracellular matrix consistent of sodium bicarbonate (1.1% final concentration; Sigma-Aldrich, #25080094), collagen I (1 mg/ml final concentration; Corning, #354249), tumor medium (DMEM high glucose (Brunschwig, #DMEM-HA) supplemented with 1 mM of sodium pyruvate (Sigma-Aldrich, S8636), 1X MEM nonessential amino acids (Gibco, #11140050), 2 mM of L-glutamine (Thermo Fisher Scientific, #25030081), 10% FBS (Brunschwig, #FBS-LE-12A), 1% penicillin–streptomycin (Gibco, #15140122) and 50 µM β-mercaptoethanol (Gibco, #11528926)), and ice-cold Cultrex UltiMatrix BME (Bio-Techne, BME001)). All components were kept ice-cold to avoid pre-mature solidification of the collagen. PDTFs were slowly thawed at 37 °C and washed in tumor medium multiple times by flushing them through a 100 µm cell strainer. One PDTF per well was placed on top of the pre-solidified matrix (15 min at 37 °C for solidification), after which a second layer of matrix was added. Plates were then incubated at 37 °C for another 15 min. After solidification, tumor medium supplemented with either anti-NKG2A (10 µg/ml, Ichorbio, ICH5017), or Isotype control (human IgG4, QA16A15: 10

µg/ml, Biolegend, #403701) was added. Ten to fifteen PDTFs were used per condition and cultures were incubated at 37 °C for 48 hours. 10 hours before harvest, a Brefeldin A (1:1000) (eBiosciences, #00-4506-51), and Monensin (1:1000) (eBiosciences, #00-4505-51) mix was added to block cytokine secretion. For flow cytometry, PDTFs were processed into single cell suspensions by enzymatic digestion, washed in PBS and filtered using a 100 µM filter mesh.

**Flow cytometry**

Tumor lesions were centrifuged at 400*g* for 5 minutes (min) in 96-well V-bottom plates. Subsequently, cells were washed with PBS, stained with fixable live/dead Zombie dyes (BioLegend) for 20 min at 4°C in PBS. After two washing steps using PBS, cells were blocked with a human Fc receptor binding inhibitor (Invitrogen) for another 20 min at 4°C in PBS. After further centrifugation, cell surface antigens using fluorophore-conjugated anti-human Abs (table S3) diluted in flow cytometry buffer (1X PBS, 0.5 mM EDTA, 2% fetal calf serum, 1% sodium azide ($NaN_3$)), for 20 min at 4°C. After two washes with FACS buffer, cells were permeabilized and fixed for intranuclear staining using Foxp3/Transcription Factor Staining Buffer Set (Invitrogen, #00-5523-00) and then stained intranuclearly with antibody cocktails in 1X Permeabilization buffer (Invitrogen, #00-8333-56). After two washes with Permeabilization buffer, the samples were resuspended in flow cytometry buffer and acquired using Aurora (Cytek). FlowJo (V.10.10.0) was used for analysis.

**Transduction of tumor cells**

To generate an HLA-E expressing A549-b2m$^{ko}$ (RRID: CVCL_C3CC) cell line, B2M HLA-E lentiviral particles (BPS Bioscience) were added at a multiplicity of infection (MOI) of 8 for 48 hours in tumor cell culture medium (DMEM high glucose (Brunschwig, #DMEM-HA) supplemented with 1X MEM nonessential amino acids (Gibco, #11140050), 2 mM HEPES (Gibco, #15630080), 10% heat-inactivated fetal bovine serum (FBS) (Brunschwig, #FBS-LE-12A), 1% penicillin–streptomycin (Gibco, #15140122) and 50 µM β-mercaptoethanol (Gibco, #11528926)). Subsequently, cells were expanded every 2 days with fresh medium supplemented with 0.5 µg/ml puromycin (Gibco, #A1113802).

**Ex vivo expansion**

Thawed tumor lesions were rested overnight in NK cell culture medium supplemented with 10 ng/ml IL-15. Subsequently, cells were cultured for 7 days under three conditions: 10 ng/ml IL-15 alone, irradiated K562 (RRID: CVCL_0004; 90 Gy; 2:1 E:T ratio), or a combination of both. Media and IL-15 were replenished every 48 hours. Following phenotypic and quantitative assessment at day 0 and day 7, viable (Zombie $NIR^-$) $CD3^-CD56^+$ NK cells were sorted into $CD103^+CD49a^+$ and $CD103^-CD49a^-$ subsets using Cytek Aurora Cell Sorter. Sorted populations were then utilized in functional killing assays against A549-$b2m^{ko}$ target cells, as described above.

**Bulk RNA sequencing**

In vitro-generated tumor-associated NK cells of four donors were sorted for $CD103^+CD49a^+$ and $CD103^-CD49a^-$ populations as described above. Cell pellets were subsequently be stored at -80°C until further use. RNA extraction was performed using the Qiagen RNeasy Plus Mini Kit (250) (Qiagen, #74136) according to manufacturer's instructions. Quality control, including length profiling and concentration determination (Agilent Fragment Analyzer), and library preparation (SMART-Seq Total RNA-Seq Single Cell [ZapR Mammalian], Takara) were conducted by the Genomics Core Facility at the University of Basel. Sequencing was performed on the Element Biosciences AVITI platform, generating 75 nt paired-end reads. Raw FASTQ files were processed using the nf-core/rnaseq pipeline (v3.15.1) (doi: 10.5281/zenodo.1400710) *(125)*, aligning reads to the Ensembl human hg38 genome (release 115). Read quantification was performed via the STAR/Salmon workflow following UMI deduplication. Subsequent statistical analyses were conducted in R (v4.4.2). Differential expression analysis was performed using DESeq2 (v1.46.0) *(126)*, incorporating donor origin as a covariate to account for inter-individual variability. Genes with fewer than 10 counts were excluded. To refine fold-change estimates for lowly expressed genes, the apeglm shrinkage method was applied. Functional enrichment was assessed via GSEA using clusterProfiler (v4.4.2) *(121)*.

**Re-analysis of published datasets with response information to ICB**

Published bulkRNA and scRNA-seq data of cancer patients with information about response to ICB were downloaded (bulkRNA: Jung *et al. (81)* (GSE135222), Chen *et al. (127)* (GSE236581), Bassez *et al. (128)* (http://biokey.lambrechtslab.org/)). Analyses of single cell datasets were performed using the standard Seurat workflow, NK cells were extracted based on published dataset annotations and enrichment of taNK cells (top 30 genes) per patient was performed using the UCell and escape v1.12.0 *(120)* package. The NK matrix was deconvoluted from the bulk RNA-seq datasets as mentioned above using the BayesPrism package. The maxRank was increased to 2000 genes for the enrichment analysis of bulkRNA datasets.

**Integration with non-tumor lung**

Non-tumor lung datasets were obtained from Sikkema *et al. (62)* (data available at cellxgene) and Bischoff *et al. (63)* (data available at code ocean capsule). For Sikkema *et al.*, only normal lung tissue was selected and NK cells were extracted from the dataset using the dataset annotation. For Bischoff *et al.*, normal lung tissue datasets were downloaded, the dataset processed using the standard Seurat workflow and NK cells were subsetted based clusters expressing *NCAM1, NKG7* and *GNLY*, while negative for *CD3D* and *CD3G.* Subsequently, the RNA dataset of our lung tumor NK cells was extracted, all datasets merged, the count matrices for each donor and dataset log-normalized, scaled, and CCA integrated as mentioned above. 15 PCs were used for clustering and dimensional reduction and clusters were annotated using DEG analysis and signature scoring as mentioned above.

**Inference of genes and eRegulons evolution along the trajectory**

To validate the identified transcriptional trajectories and gain deeper insight into the transition from NK3_GZMK to NK3_ENTPD1, we performed pseudotime analysis of these populations using Monocle3 *(129)* (v1.3.1) on all samples combined. The *learn_graph* function was run with the parameter 'ncenter = 100' to prevent over-branching of the trajectory. The starting point was defined as the terminal branch within the NK3_GZMK population, based on MultiVelocity and diffusion map analyses. Pseudotime was then computed using the *order_cells* function.
To identify genes and regulons correlated with pseudotime progression along the principal graph, we performed Moran's I test. The top 100 genes with a q-value < 0.05 and the highest Moran's I scores were selected and visualized along pseudotime (z-scored expression) using the *Heatmap* function from the ComplexHeatmap library (v2.6.2). For regulon analysis, we focused specifically on activators (i.e., those associated with increased chromatin accessibility and up-regulation of target gene expression).

**Overall survival analysis**

Expression data of TCGA and clinical information for LUAD and LUSC were downloaded using the TCGAbiolinks *(130)* v2.32.0. NK expression matrix was deconvoluted by BayesPrism *(131)* v.2.2.2 using the single cell dataset from Kim *et al. (132)* (GSE131907) as reference for NSCLC and the single cell dataset from Bassez *et al. (128)* (http://biokey.lambrechtslab.org/) for breast cancer to score the signature genes (top 10) identified in taNK cells. The patients were assigned as high (top 25%) and low (bottom 25%). The Cox proportional hazards model was used implemented in the R package survival *(133)* v3.7-0. Next, the survival curves were generated using the Kaplan–Meier formula. For plotting, the *ggsurvplot* function of the survminer *(134)* v.0.4.9 package was used.

**Diffusion map analysis**

Diffusion-map algorithms implemented in the R package destiny *(85)* (v3.4.0) were used to infer pseudotime. The analysis was performed separately on RNA expression (Fig. 5A and 5B) and GeneActivity scores inferred from ATAC data (fig. S11C and S11D) in NK3 cells pooled from all samples. For the RNA analysis, diffusion map embedding was performed using the variable genes identified during NK3 preprocessing. For GeneActivity, the analysis was conducted on the top 5% most frequently shared features across cells. To prevent individual batch effect (at the sample level), the *RunFastMNN* function implemented in the R package batchelor *(135)* (v1.6.3) was used. The corrected expression matrix was then used as input to generate diffusion maps using the *DiffusionMap* function, with the parameters set to censor_val = 30 and censor_range = c(30, 40). The Destiny algorithm automatically identified three 'root' cells. For the ATAC based Gene Activity, the first diffusion component was removed as it was correlated with quality of sequencing and therefore noised the diffusion analysis. The main root was selected from the NK3_GZMK cluster, at the beginning of the directed streamline inferred by MultiVelocity. Diffusion pseudotime for all cells was then calculated using the *DPT* function.

**Spatial Transcriptomics**

<u>Sample selection</u>

Samples were obtained from a male, non-smoking patient diagnosed with primary, treatment-naïve NSCLC (adenocarcinoma, table S2). Multiple anatomical regions were included: tumor core, tumor invasive front, and adjacent lung tissue. Tumor classification, region annotation and TMA construction were performed by the Department of Pathology, Hannover Medical School (MHH).

Tissue microarray (TMA) construction

TMAs were generated from formalin-fixed, paraffin-embedded (FFPE) tissue. 4 × 4 mm cores of regions of interest were transferred into TMA blocks. Blocks were sectioned at 5 µm and mounted onto a Xenium slide (10x Genomics; CG000578, Rev. A). Adjacent sections were stained with hematoxylin and eosin (H&E) for histomorphological evaluation.

Xenium in situ gene expression workflow and on-board analysis

In situ gene expression profiling was performed using the Xenium In Situ Gene Expression platform (10x Genomics) according to the manufacturer's protocol (CG000749, Rev. B). The predesigned Human Immuno-Oncology Panel (380 genes) supplemented with a custom add-on panel (100 genes; FTT2GF) was used. Antibody-based cell segmentation staining was performed based on membrane boundary, intracellular RNA and nuclear staining. Slides were processed on the Xenium Analyzer using Xenium software v4.0, with instrument setup, region selection and run initiation performed according to the Xenium Analyzer User Guide (CG000584, Rev. K).

Xenium Analysis Pipeline

Xenium output files were primarily processed using R (v.4.5.0), Seurat (v.5.2.1) and arrow (v.18.1.0.1). Initial quality control parameters were guided by the Xenium Onboard Analyzer (v.4.0.1.0). Cells with fewer than 20 detected features (nFeatures < 20) or fewer than 50 transcript counts (nCount < 50) were excluded from downstream analysis. Data were normalized using SCTransform followed by dataset integration via reciprocal PCA. Clustering was performed using the Seurat pipeline and visualized as UMAP. Cluster annotation was performed based on DEGs calculated with the *FindAllMarkers* function. Canonical marker genes were used to identify cell populations, including T cells (*CD3D, TRAC, FOXP3, CTLA4*), B cells (*MS4A1, MZB1, CD79A*), monocytes (*MARCO, LYZ*), epithelial and endothelial cells (*EPCAM, KRT7, KRT5, FOXJ1, PECAM1, VWF, ACKR1, PROX1*), and NK cells (*NKG7, GNLY, GZMB*). Automated annotation using SingleR (v.2.13.1) was additionally performed to support manual cluster classification. To distinguish NK cells from $CD8^+$ T cells, module scoring was performed using *AddModuleScore_UCell* from the UCell package *(118)*. A *CD3D*-based signature score threshold of 0.55 was applied. Cells exceeding this threshold were classified as "T-like" and removed from the NK cell compartment, whereas remaining cells were defined as "NK-cleaned". Further subclassification of NK cells into taNKs was based on the taNK gene signature derived from Multiome data. Of the signature genes, 31 were represented in the Xenium panel and used for

UCell scoring. Cells with a signature score > 0.61 were classified as taNKs, while cells below this threshold were defined as non-taNKs.

Visualization

Spatial visualization was performed using Xenium Explorer (v.4.1.0). Cluster identities and annotations generated in R were exported as .csv files and imported into Xenium Explorer for spatial mapping. To identify *ENTPD1* (CD39)-positive taNK cells, expression of *ENTPD1, ITGAE, ITGA1, NKG*7, and *CD3D* was visualized. Images were exported at a resolution of 5100 × 2925 pixels and 600 DPI.

**Visualizations**

Heatmaps were plotted using the R package ComplexHeatmap *(136)* v2.20.0, volcano plots with the R package EnhancedVolcano *(137)* v1.22.0 and boxplots using ggplot2 v3.5.1. Furthermore, the package *DittoSeq (138)* was used. Schematic workflows were illustrated using Biorender.com. Flow cytometry plots were generated using GraphPad Prism v10.3.0 and FlowJo v10.10.0.

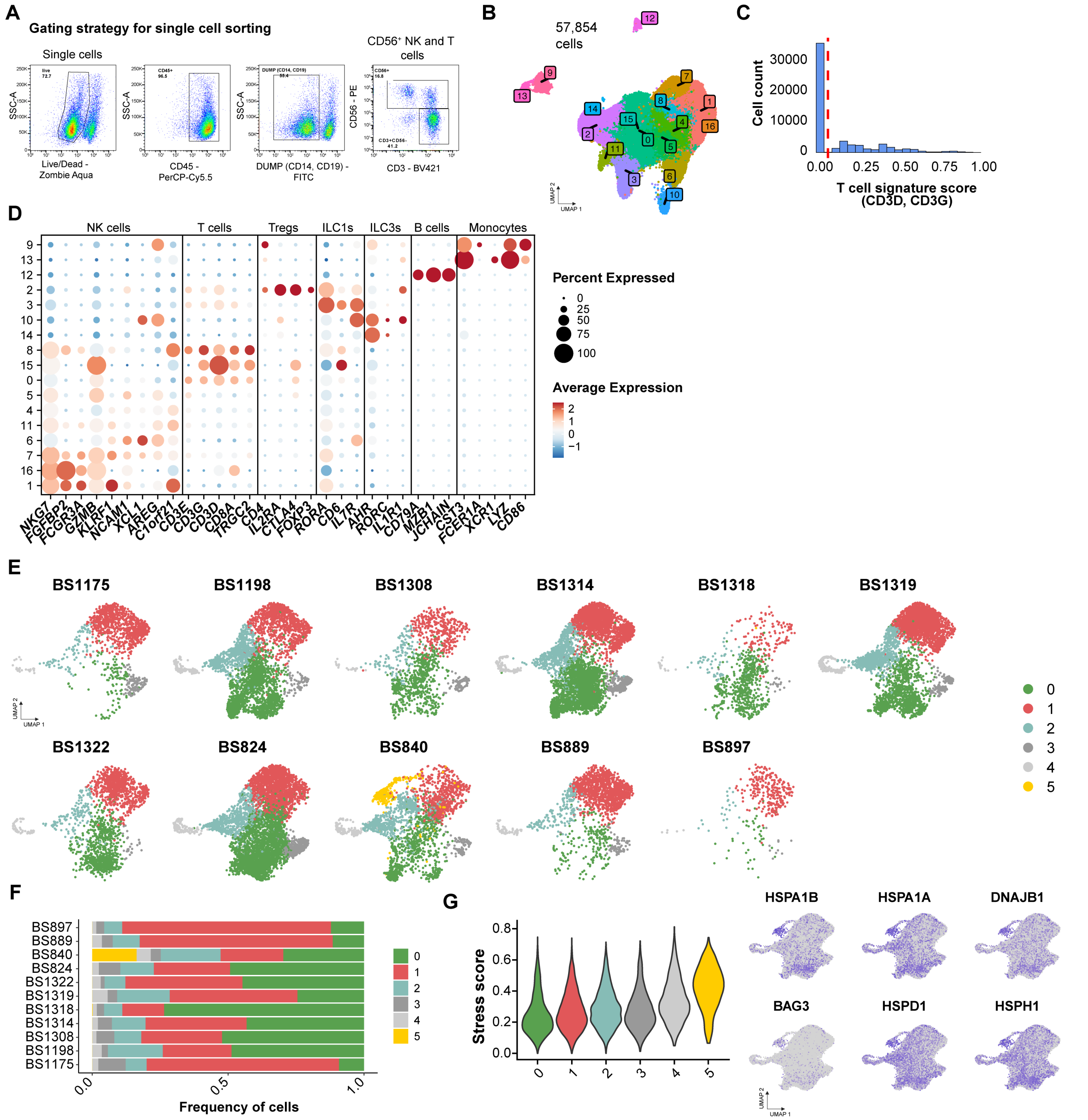


**Fig. S1 Technical NK cells extraction and intratumoral heterogeneity of lung tumor NK cells**

**(A)** Gating strategy for sorting. Sorted cells were used as input for the single cell experiment. **(B)** Weighted nearest neighbor (WNN) integrated Uniform Manifold Approximation and Projection (UMAP) representation of all recovered cells after quality control (QC). **(C)** Histogram showing the distribution of the UCell signature score for CD3D and CD3G across all cells. Red line indicates threshold for subsetting T cells. **(D)** Average expression dot plot of indicated genes facilitating the identification of indicated cell types. **(E** and **F)** UMAP representation and bar plot of frequencies of coarse clusters split per patient. **(G)** Violin plot showing stress signature *(26)* scorings across coarse clusters and feature plots representing the expression of indicated genes.

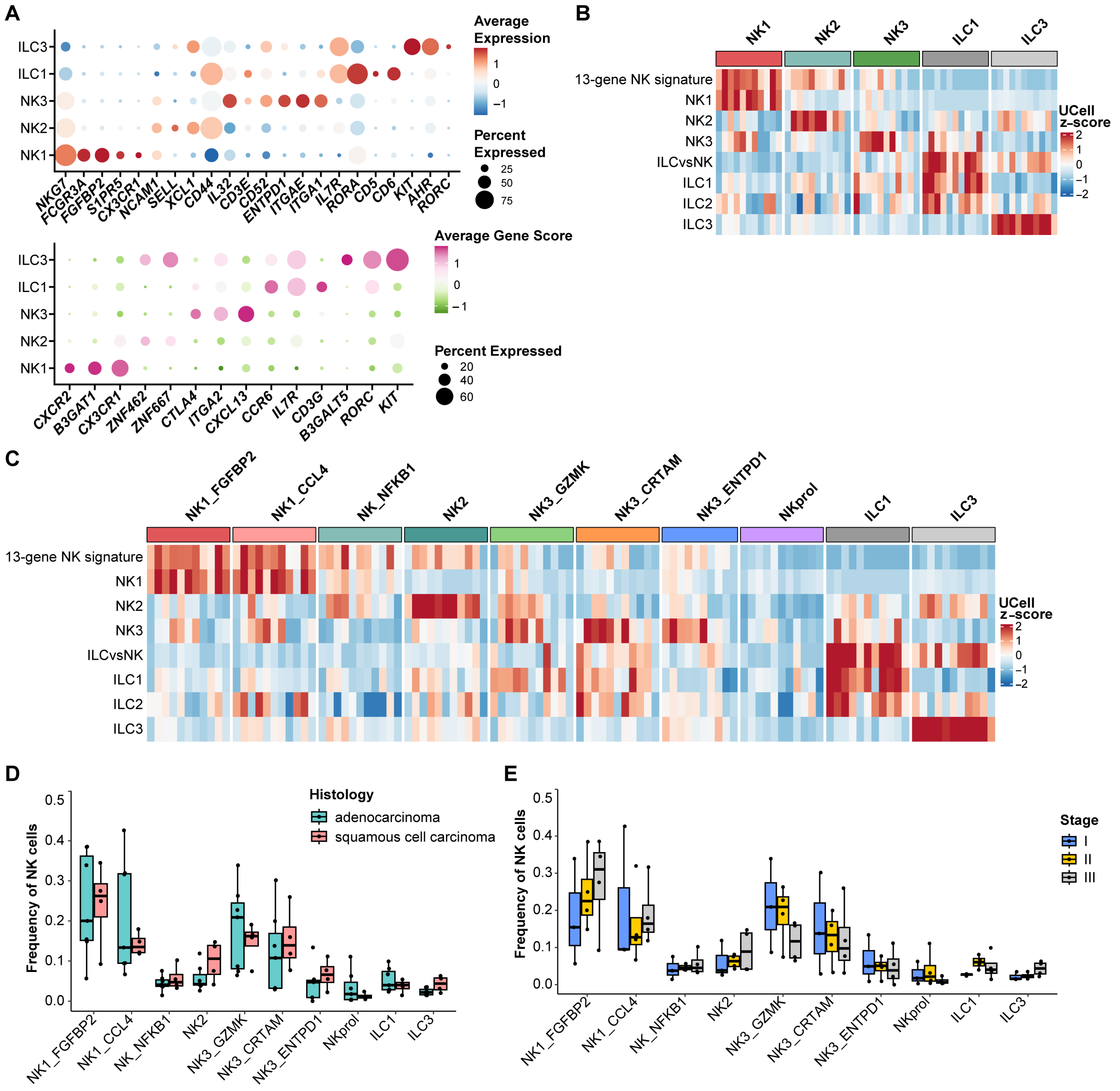


**Fig. S2 Natural killer (NK) cells from human lung cancer exhibit substantial heterogeneity**

**(A)** Average expression and gene score dotplots of indicated genes for coarse clusters identified in the combined weighted nearest neighbor (WNN) UMAP. **(B and C)** Heatmap showing row-scaled UCell signature scores of indicated signature *(11–12, 30)* across patients of (B) coarse and (C) fine cluster of the combined WNN UMAP. **(D and E)** Boxplots of NK cell frequencies within each cluster per patient comparing either tumor histology or lung cancer stage.

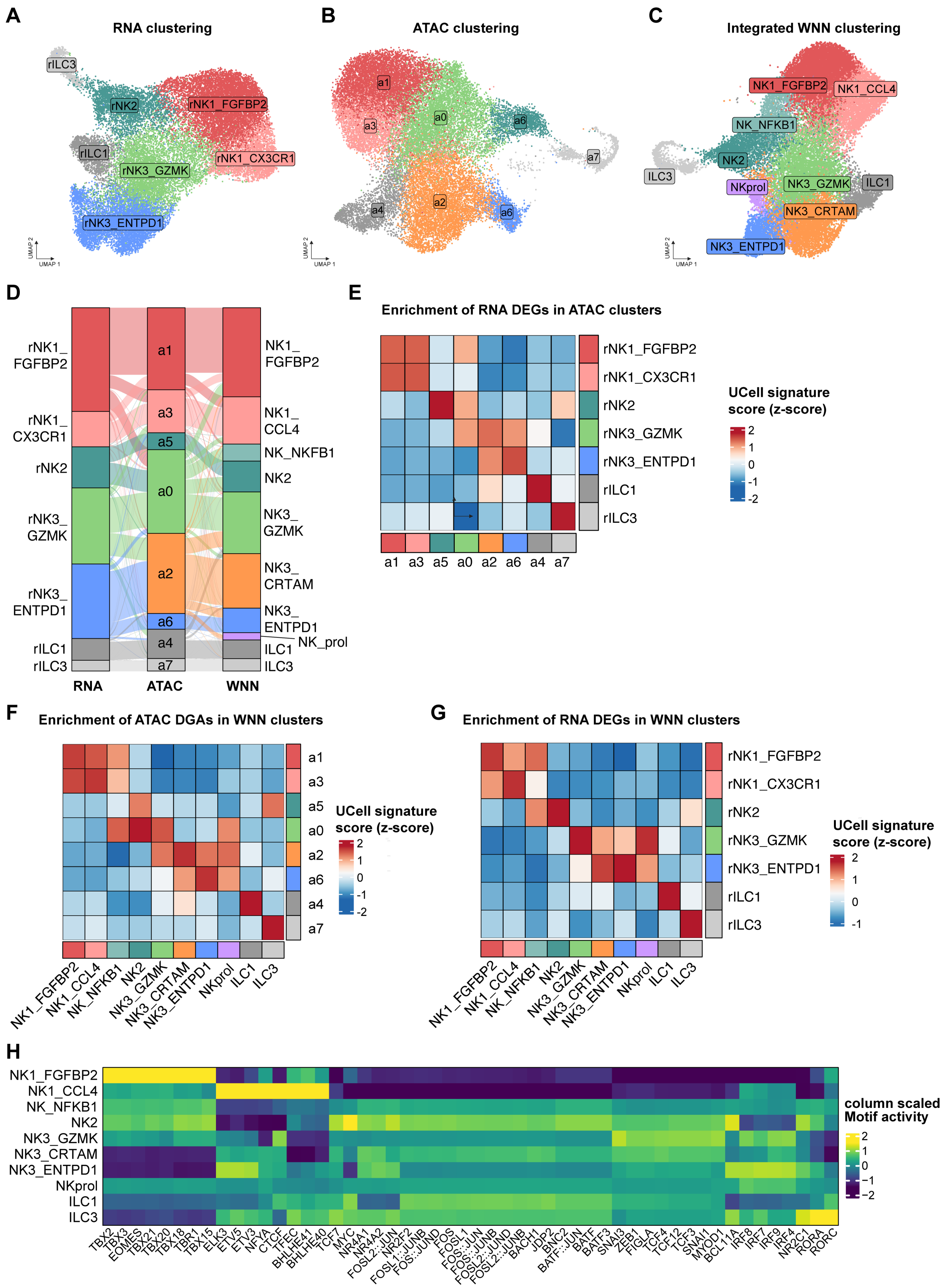
A
RNA clustering
B
ATAC clustering
C
Integrated WNN clustering
D
E
Enrichment of RNA DEGs in ATAC clusters
UCell signature score (z-score)
F
Enrichment of ATAC DGAs in WNN clusters
G
Enrichment of RNA DEGs in WNN clusters
H
column scaled Motif activity

248

**Fig. S3 Comparison of Single-Modality and Integrated (WNN) Clustering**

(**A** to **C**) UMAP visualizations of NK cell subsets clustered using (A) single nucleus RNA-sequencing data alone, (B) single nucleus ATAC-sequencing data alone, and (C) the WNN integrated analysis (as used in the main manuscript). (**D**) Alluvial plot showing the flow of single-cell assignments across the three clustering modalities (RNA → ATAC → WNN). (**E** to **G**) Heatmaps depicting average UCell signature score (z-score): per ATAC cluster of RNA cluster DEGs (top 50), using (E) gene activity, per WNN cluster of ATAC cluster differential gene activities (DGAs), using (F) RNA expression and per WNN cluster of RNA cluster DEGs (top 50), using (G) RNA expression. (**H**) Heatmap showing average transcription factor (TF) motif activity across different NK cell subtypes.

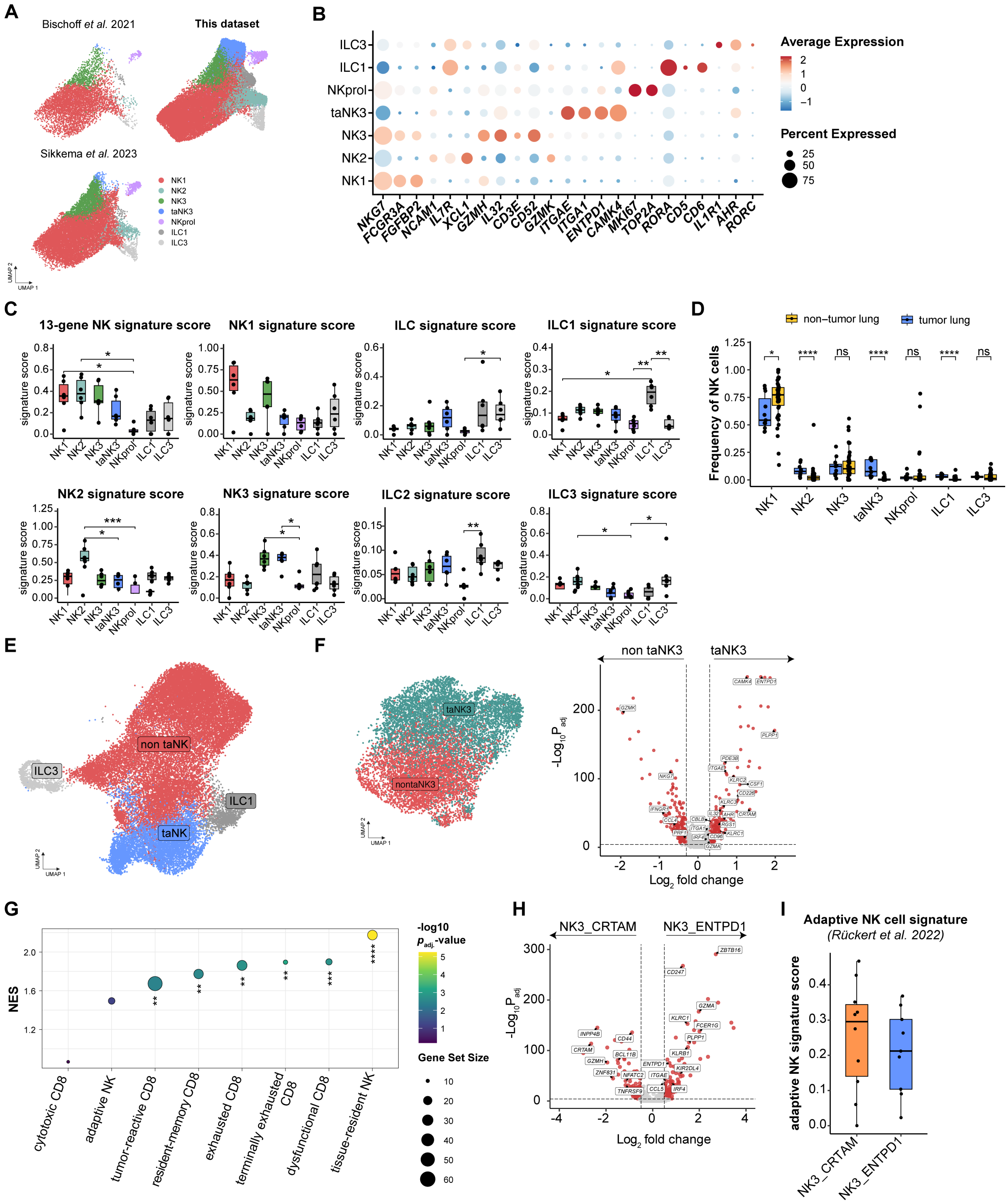
A
Bischoff et al. 2021
This dataset
Sikkema et al. 2023
NK1
NK2
NK3
taNK3
NKprol
ILC1
ILC3
UMAP 2
UMAP 1
B
ILC3
ILC1
NKprol
taNK3
NK3
NK2
NK1
NKG7
FCGR3A
FGFBP2
NCAM1
IL7R
XCL1
GZMH
IL32
CD3E
CD52
GZMK
ITGAE
ITGA1
ENTPD1
CAMK4
MKI67
TOP2A
RORA
CD5
CD6
IL1R1
AHR
RORC
Average Expression
Percent Expressed
C
13-gene NK signature score
NK1 signature score
ILC signature score
ILC1 signature score
NK2 signature score
NK3 signature score
ILC2 signature score
ILC3 signature score
signature score
D
non−tumor lung
tumor lung
Frequency of NK cells
ns
E
non taNK
taNK
ILC3
ILC1
F
taNK3
nontaNK3
non taNK3
Log2 fold change
G
NES
-log10 Padj.-value
Gene Set Size
cytotoxic CD8
adaptive NK
tumor-reactive CD8
resident-memory CD8
exhausted CD8
terminally exhausted CD8
dysfunctional CD8
tissue-resident NK
H
NK3_CRTAM
NK3_ENTPD1
Log2 fold change
I
Adaptive NK cell signature
(Rückert et al. 2022)
adaptive NK signature score

**Fig. S4 Comparison of non-tumor-associated NK cells with tumor-associated NK cells**

**(A)** UMAP representations of combined non-tumor and tumor lung NK cells split by dataset (50,088 cells). **(B)** Average expression dotplots of indicated genes for clusters identified in the combined non-tumor and tumor lung UMAP. **(C)** Boxplots showing Ucell signature scores of indicated signatures *(11–12, 30)*. Each dot represents a study. (n = 11) **(D)** Boxplot showing the cell frequencies comparing non-tumor lung and tumor lung NK cells within each cluster. Each dot represents one donor. (tumor lung: n = 11, non-tumor lung: n = 50) **(E)** UMAP representation of clusters summarized as non-tumor-associated and tumor-associated NK cells. **(F)** UMAP representation of NK3 only clusters summarized as non-tumor-associated and tumor-associated NK cells (left). Volcano plot showing differentially expressed genes (DEG) between the taNK3 and non taNK3 clusters (right). **(G)** Dot plot showing normalized gene set enrichment analysis (GSEA) scores (NES) of published signatures *(13, 31, 54, 65–66, 119)* in tumor-associated NK3 cells signature. **(H)** Volcano plot showing differentially expressed genes (DEG) between the NK3_ENTPD1 and NK3_CRTAM clusters. **(I)** Boxplots showing Ucell signature scores between NK3_CRTAM and NK3_ENTPD1 of the adaptive NK cell signature. Each dot represents one donor. Data were analyzed using (C) non-parametric Kruskal-Wallis test with post-hoc Dunn's multiple comparison, adjusted P value, (D) Wilcoxon signed-rank test, (F and H) the MAST hurdle model and *P*-values were calculated using a likelihood ratio test and adjusted for multiple comparisons using Bonferroni correction, (G) permutation-based testing with adjustment for multiple comparisons using ClusterProfiler.

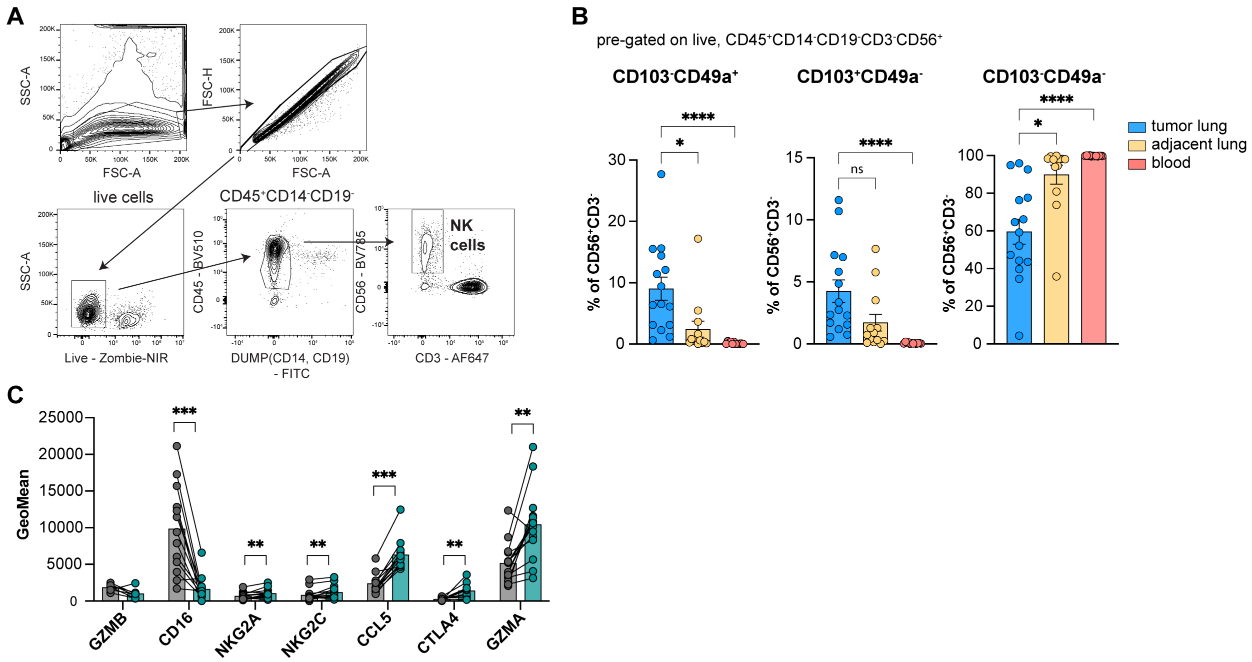


**Fig. S5. Multiplexed flow cytometry validates the identity of tumor-associated NK cells.**

**(A)** Gating strategy for identifying NK cells. **(B)** Barplot (Mean ± SEM) of indicated NK cell populations in tumor lung and adjacent lung quantified using flow cytometry. (tumor lung: n=15, adjacent lung: n=13, blood: n=12) **(C)** GeoMean of indicated proteins comparing $CD103^{+}CD49^{+}$ NK cells (petrol) and $CD103^{-}CD49^{-}$ NK cells (grey) from baseline tumor lung digests (n = 14). Data were analyzed by (B) Kruskal-Wallis test with post-hoc Dunn's multiple comparisons, (C) paired wilcoxon signed-rank test.

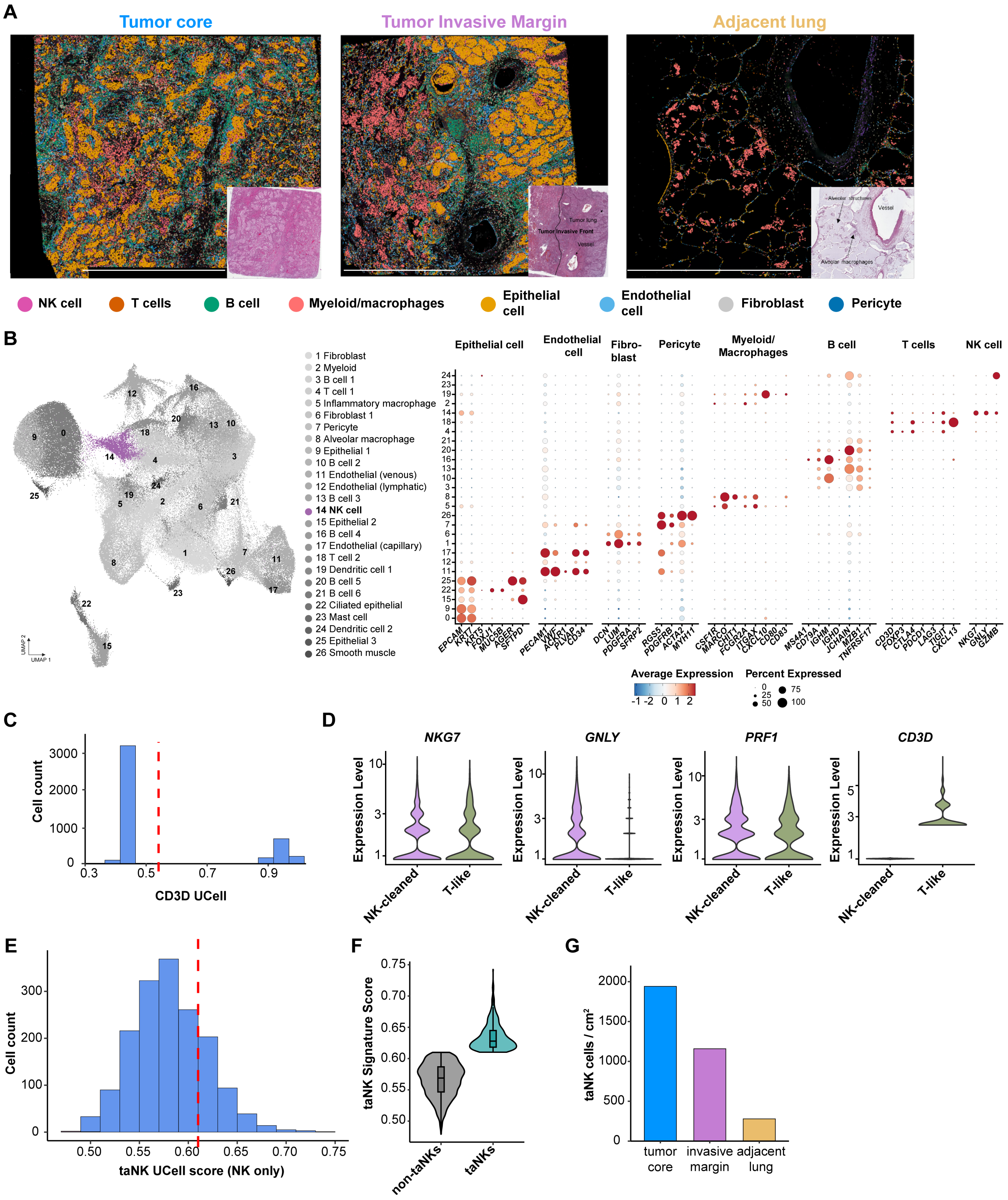
A
Tumor core
Tumor Invasive Margin
Adjacent lung
NK cell
T cells
B cell
Myeloid/macrophages
Epithelial cell
Endothelial cell
Fibroblast
Pericyte
B
1 Fibroblast
2 Myeloid
3 B cell 1
4 T cell 1
5 Inflammatory macrophage
6 Fibroblast 1
7 Pericyte
8 Alveolar macrophage
9 Epithelial 1
10 B cell 2
11 Endothelial (venous)
12 Endothelial (lymphatic)
13 B cell 3
14 NK cell
15 Epithelial 2
16 B cell 4
17 Endothelial (capillary)
18 T cell 2
19 Dendritic cell 1
20 B cell 5
21 B cell 6
22 Ciliated epithelial
23 Mast cell
24 Dendritic cell 2
25 Epithelial 3
26 Smooth muscle
Epithelial cell
Endothelial cell
Fibro-blast
Pericyte
Myeloid/ Macrophages
B cell
T cells
NK cell
EPCAM KRT7 KRT5 FOXJ1 MUC5B AGER SFTPD
PECAM1 VWF ACKR1 PLVAP CD34
DCN LUM PDGFRA SFRP2
RGS5 PDGFRB ACTA2 MYH11
CSF1R MARCO CHIT1 FCGR2A ITGAX CXCL10 CD80 CD83
MS4A1 CD79A IGHM IGHD JCHAIN MZB1 TNFRSF17
CD3D FOXP3 CTLA4 PDCD1 LAG3 TIGIT CXCL13
NKG7 GNLY GZMB
Average Expression
Percent Expressed
C
Cell count
CD3D UCell
D
NKG7
GNLY
PRF1
CD3D
Expression Level
NK-cleaned
T-like
E
Cell count
taNK UCell score (NK only)
F
taNK Signature Score
non-taNKs
taNKs
G
taNK cells / cm²
tumor core
invasive margin
adjacent lung

**Fig. S6. Xenium spatial transcriptomics workflow and validation of tumor-associated NK cell identity in situ.**
**(A)** Spatial mapping of unbiased Xenium-derived cell clusters across the tumor core, invasive margin, and adjacent lung of a lung adenocarcinoma (LUAD) patient, with corresponding H&E-stained sections provided as insets for anatomical context. Scale Bar: 2 mm. **(B)** UMAP representation of 26 identified cell clusters (left) and a dot plot (right) validating cell type annotations using canonical marker genes for epithelial, stromal, and immune lineages. **(C)** Distribution of the *CD3D* UCell signature score within the NK cell cluster (cluster 14), where a threshold (red dashed line, < 0.55) was applied to isolate a "cleaned" NK cell population from T-cell contamination. **(D)** Expression levels of *NKG7*, *GNLY*, *PRF1*, and *CD3D* in the "NK-cleaned" subset compared to the "T-like" ($CD3D^{high}$) population partitioned by the threshold in **(C)**. **(E)** Distribution of the taNK signature score (using top 30) within the cleaned NK cell population across all tissues, with a threshold (red dashed line, > 0.61) used to define the taNK subset. **(F)** Enrichment of the taNK signature score in the taNK-high population compared to the taNK-low population as defined by the threshold in **(E)**. **(G)** Quantitative density ($cells/cm^2$) of annotated taNKs across the tumor core, invasive margin, and adjacent lung tissue (n = 1).

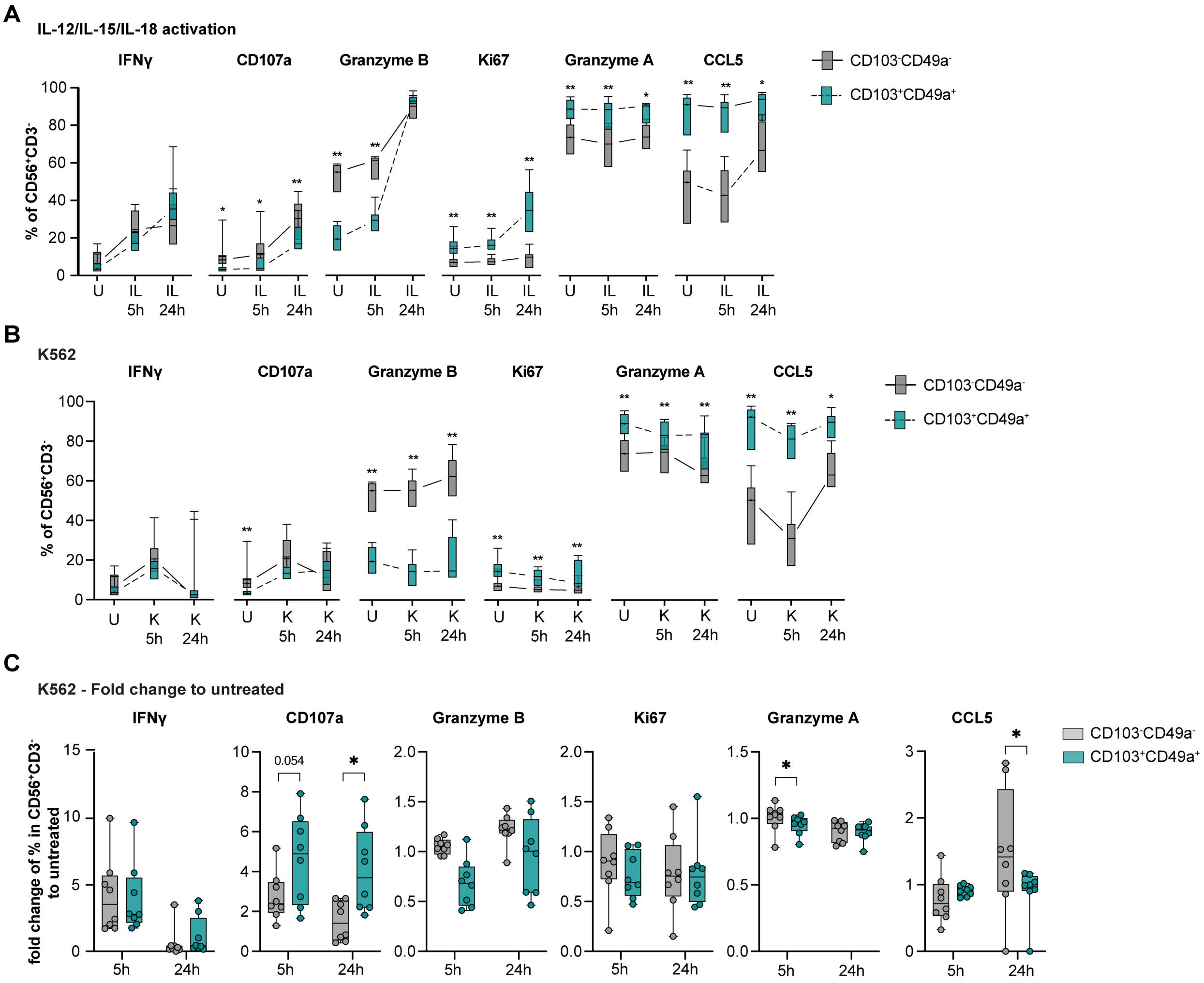


**Fig. S7: Restimulation of tumor-associated lung NK cells reveals cytotoxic potential. (A and B)** Percentage of indicated markers in CD103$^+$CD49a$^+$ (petrol) and CD103$^-$CD49a$^-$ NK cells (grey) from lung tumors upon IL-12/IL-15/IL-18 (IL, in A) treatment and in a co-culture with K562 (K, in B) for 5 or 24 hours and untreated (U) for 5 hours. (5 h: n = 8, 24 h: n = 8). **(C)** Fold change in percentage of indicated markers in CD103$^+$CD49a$^+$ (petrol) and CD103$^-$CD49a$^-$ NK cells (grey), relative to untreated, from lung tumors in a co-culture with K562 for 5 or 24 hours. (5 h: n = 8, 24 h: n = 8). Statistical significance was determined using paired Wilcoxon signed-rank test. **P < 0.05, **P < 0.01, ***P < 0.001, ****P < 0.0001*

291

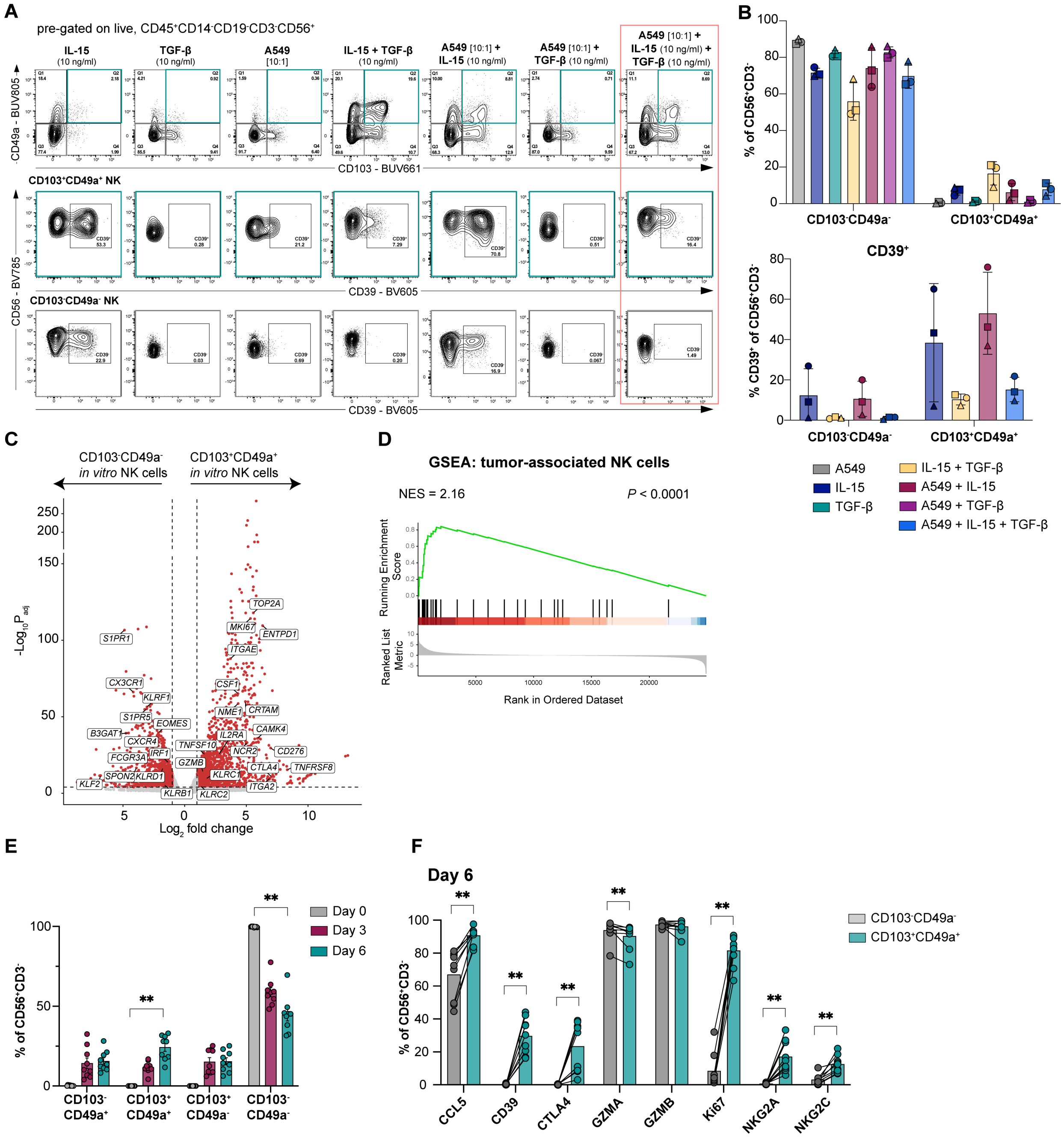
A
pre-gated on live, CD45+CD14-CD19-CD3-CD56+
IL-15 (10 ng/ml)
TGF-β (10 ng/ml)
A549 [10:1]
IL-15 + TGF-β (10 ng/ml)
A549 [10:1] + IL-15 (10 ng/ml)
A549 [10:1] + TGF-β (10 ng/ml)
A549 [10:1] + IL-15 (10 ng/ml) + TGF-β (10 ng/ml)
CD49a - BUV805
CD103 - BUV661
CD103+CD49a+ NK
CD103-CD49a- NK
CD56 - BV785
CD39 - BV605
B
% of CD56+CD3-
CD103-CD49a-
CD103+CD49a+
CD39+
% CD39+ of CD56+CD3-
A549
IL-15
TGF-β
IL-15 + TGF-β
A549 + IL-15
A549 + TGF-β
A549 + IL-15 + TGF-β
C
CD103-CD49a- in vitro NK cells
CD103+CD49a+ in vitro NK cells
-Log10 Padj
Log2 fold change
TOP2A
MKI67
ENTPD1
ITGAE
S1PR1
CX3CR1
KLRF1
CSF1
S1PR5
EOMES
NME1
CRTAM
B3GAT1
CXCR4
IL2RA
CAMK4
IRF1
TNFSF10
NCR2
CD276
FCGR3A
GZMB
CTLA4
TNFRSF8
SPON2
KLRD1
KLRC1
KLF2
ITGA2
KLRB1
KLRC2
D
GSEA: tumor-associated NK cells
NES = 2.16
P < 0.0001
Running Enrichment Score
Ranked List Metric
Rank in Ordered Dataset
E
% of CD56+CD3-
Day 0
Day 3
Day 6
CD103- CD49a+
CD103+ CD49a+
CD103+ CD49a-
CD103- CD49a-
F
Day 6
% of CD56+CD3-
CD103-CD49a-
CD103+CD49a+
CCL5
CD39
CTLA4
GZMA
GZMB
Ki67
NKG2A
NKG2C

**Fig. S8: Validation of the *in vitro* tumor-associated NK cell platform. (A** and **B)** Representative (A) contour plots and (B) quantitative bar plots (Mean ± SD) showing the induction of CD103 and CD49a on isolated healthy-donor (HD) NK cells, and subsequent CD39 expression within the resulting subsets, after 3 days of culture under indicated stimulations: IL-15 (10 ng/ml), TGF-β (10 ng/ml), irradiated A549 (E:T, 10:1), IL-15 (10 ng/ml) + TGF-β (10 ng/ml), A549 (E:T, 10:1) + IL-15 (10 ng/ml), A549 (E:T, 10:1) + TGF-β (10 ng/ml), and A549 (E:T, 10:1) + IL-15 (10 ng/ml) + TGF-β (10 ng/ml). (n = 3) **(C)** Volcano plot displaying differential expressed genes (DEGs) from bulk RNA sequencing comparing sorted CD103$^-$CD49a$^-$ or CD103$^+$CD49a$^+$ NK cells after 6 days in the taNK platform exposed to A549 (E:T, 10:1) + IL-15 (10 ng/ml) + TGF-β (10 ng/ml). (n = 4) **(D)** Gene signature enrichment (GSEA) of top 30 gene signature of tumor-associated lung NK cells from the Multiome dataset against the CD103$^+$CD49a$^+$ *in vitro* taNK cell signature. (n = 4) **(E)** Kinetic analysis of NK cell subset distribution (CD103/CD49a) within the taNK platform over 0, 3, and 6 days of culture (n=9). **(F)** Bar plots showing the percentage of indicated proteins comparing CD103$^+$CD49a$^+$ NK cells (petrol) and CD103$^-$CD49a$^-$ NK cells (grey) from taNK platform on day 6. (n = 9). Data were analyzed by (E) non-parametric Friedman test with post-hoc Dunn's multiple comparisons, (F) paired Wilcoxon signed-rank test. **P < 0.05, **P < 0.01, ***P < 0.001, ****P < 0.0001*

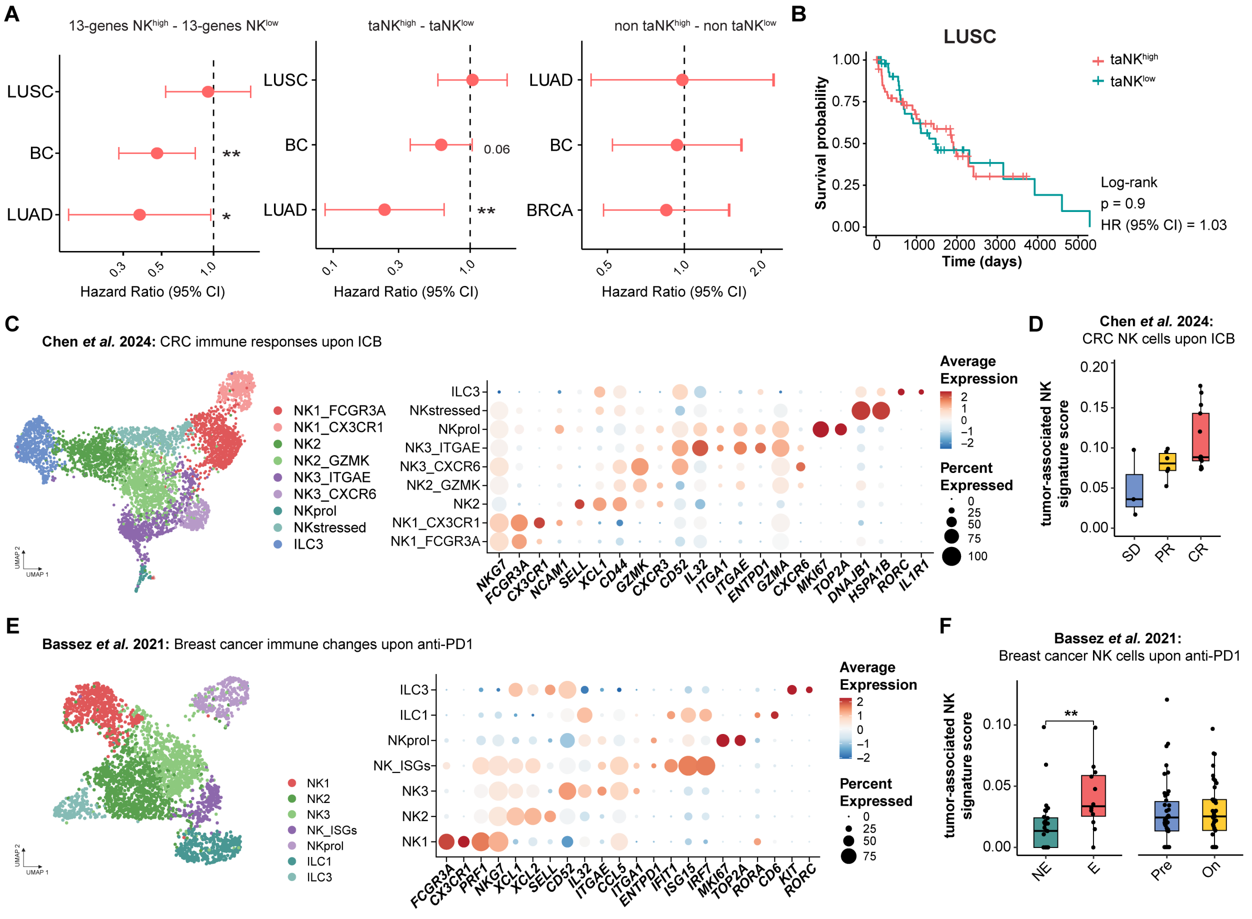


**Fig. S9: Survival analysis and reanalysis of publicly available datasets. (A)** Forest plot illustrating the impact of a 13-gene NK cell signature (*11*) (left), the tumor-associated NK signature (middle), and a non-tumor-associated NK cell signature (right) on overall survival across lung adenocarcinoma (LUAD), lung squamous cell carcinoma (LUSC), and breast cancer (BC). Hazard ratios (HR) and 95% confidence intervals (CI) are shown; y-axis is log10 transformed. **(B)** Kaplan-Maier curve showing the effect of tumor-associated NK cells on overall survival in LUSC, (ta = tumor-associated, taNK$^{high}$: top 25%, taNK$^{low}$: bottom 25%). **(C)** UMAP representation of re-analyzed intratumoral NK cells from colorectal carcinoma (CRC) upon immune checkpoint blockade (ICB) (Chen *et al.*, 2024; (*127*)) and average expression of indicated genes per cluster of re-analyzed intratumoral NK cells from CRC upon ICB (Chen *et al.*, 2024; (*127*)). **(D)** Boxplots showing UCell signature score for tumor-associated NK cells in reanalyzed single cell dataset of NK cells from CRC upon ICB (SD: stable disease, PR: partial response, CR: complete response). **(E)** UMAP representation and average expression of indicated genes per cluster of re-analyzed intratumoral NK cells from BC upon anti-PD1 treatment (Bassez *et al.*, 2021; (*128*)). **(F)** Boxplots showing UCell signature score for tumor-associated NK cells in reanalyzed single cell dataset of NK cells from breast cancer upon anti-PD1 (NE: non-expander, E: expander, Pre: Pre-treatment, On: On-treatment). Data were analyzed by (A) univariate Cox proportional hazards regression model, (B) log-rank test, (F) Wilcoxon matched-pairs signed-rank test. **P < 0.05, **P < 0.01, ***P < 0.001,*

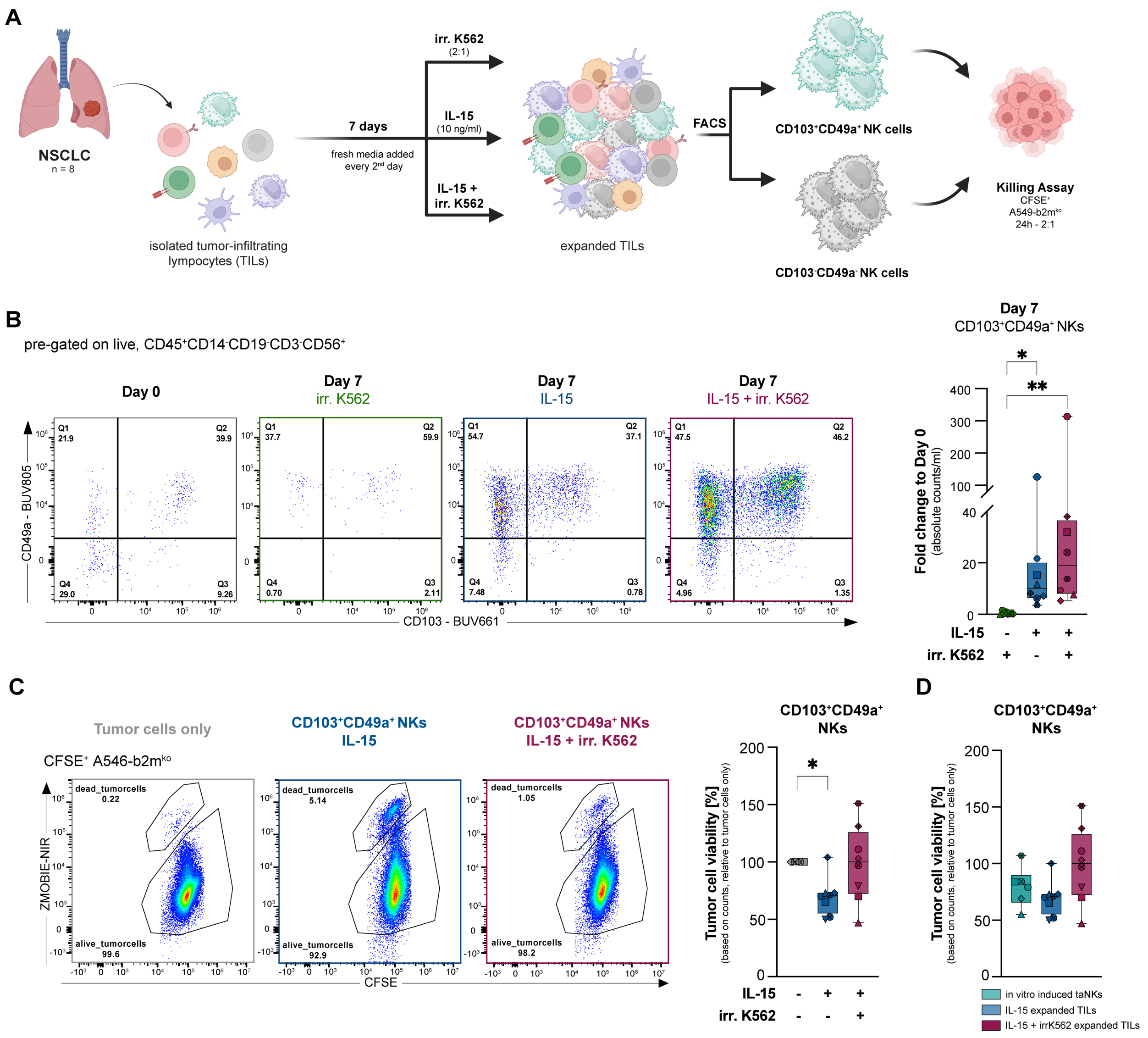


**Fig. S10: Ex vivo expansion and functional characterization of tumor-associated NKs from NSCLC. (A)** Schematic representation of ex vivo expansion assays (Graphics created with BioRender.com (*146*)). **(B)** Representative flow cytometry plots showing CD103 and CD49a expression on NK cells at day 0 and day 7 under the indicated stimulation conditions (irradiated (irr.) K562 (2:1), 10 ng/ml IL-15, or irr. K562 + IL-15) (left). Boxplot (right) showing the median fold change in absolute counts of $CD103^{+}CD49a^{+}$ NK cells relative to day 0 (n=8). **(C)** Representative plots of $ZombieNIR^{+}CFSE^{+}$ A549-b2m$^{ko}$ tumor cells after 24 hours co-culture with sorted, expanded $CD103^{+}CD49a^{+}$ NK cell (left). Tumor cell viability was determined by absolute counts and normalized to a tumor-only control (right) (n = 8) **(D)** Comparison of tumor cell viability in co-cultures with in vitro induced $CD103^{+}CD49a^{+}$ NKs (n=6) versus $CD103^{+}CD49a^{+}$ NK cells expanded from TILs with IL-15 or IL-15 + irr. K562 (n = 8) Data were analyzed using (B and C) non-parametric paired Friedman test with Dunn's multiple comparisons, (D) non-parametric Kruskal-Wallis test with Dunn's multiple comparisons. (*$*P < 0.05$, $**P < 0.01$*)

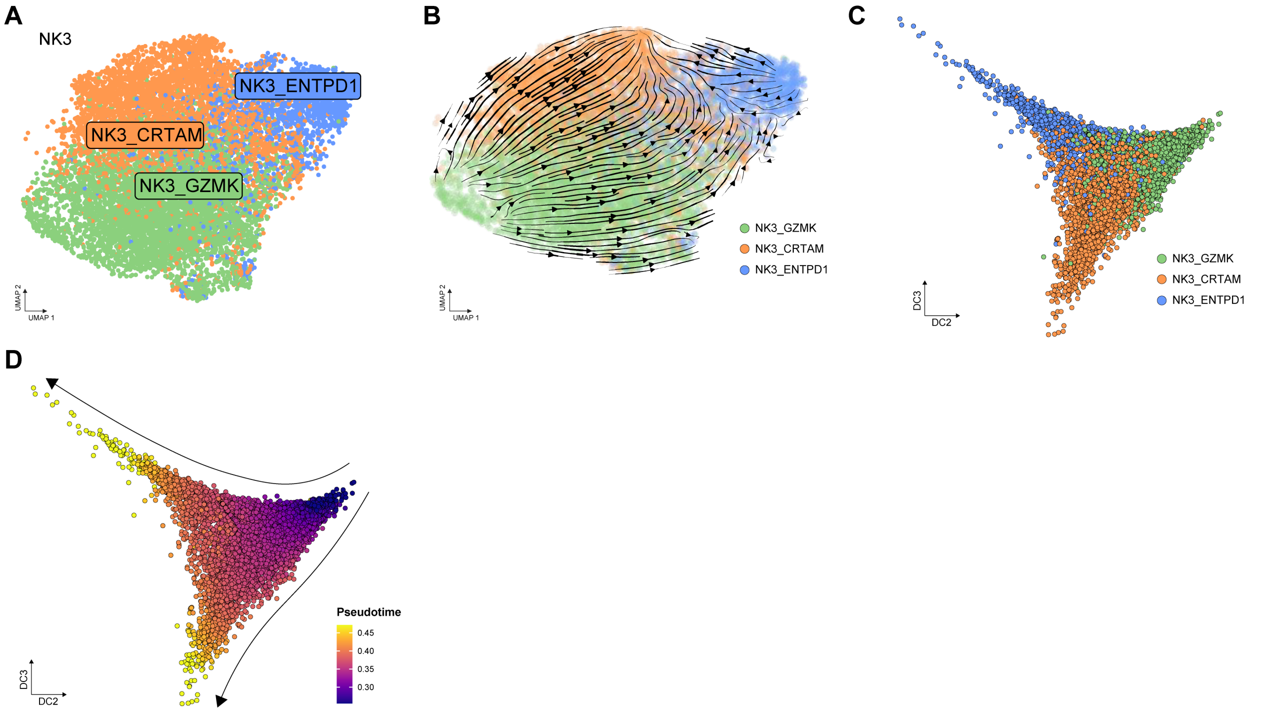


**Fig. S11: Trajectory analysis of NK3 subtypes identifies a bifurcated differentiation. (A)** UMAP of reclustered NK3 subtypes. **(B)** MultiVelocity analysis of NK3 subtypes based on combined RNA and ATAC data. **(C)** Diffusion map of single cell gene activities from NK3 cells, plotted along diffusion components DC2 and DC3. Cells colored by cluster (light green: NK3_GZMK; orange: NK3_CAMK4; blue: NK3_ENTPD1). **(D)** Pseudotime analysis of NK3 cells based on gene activity from ATAC dataset illustrated using the diffusion-map approach.

299

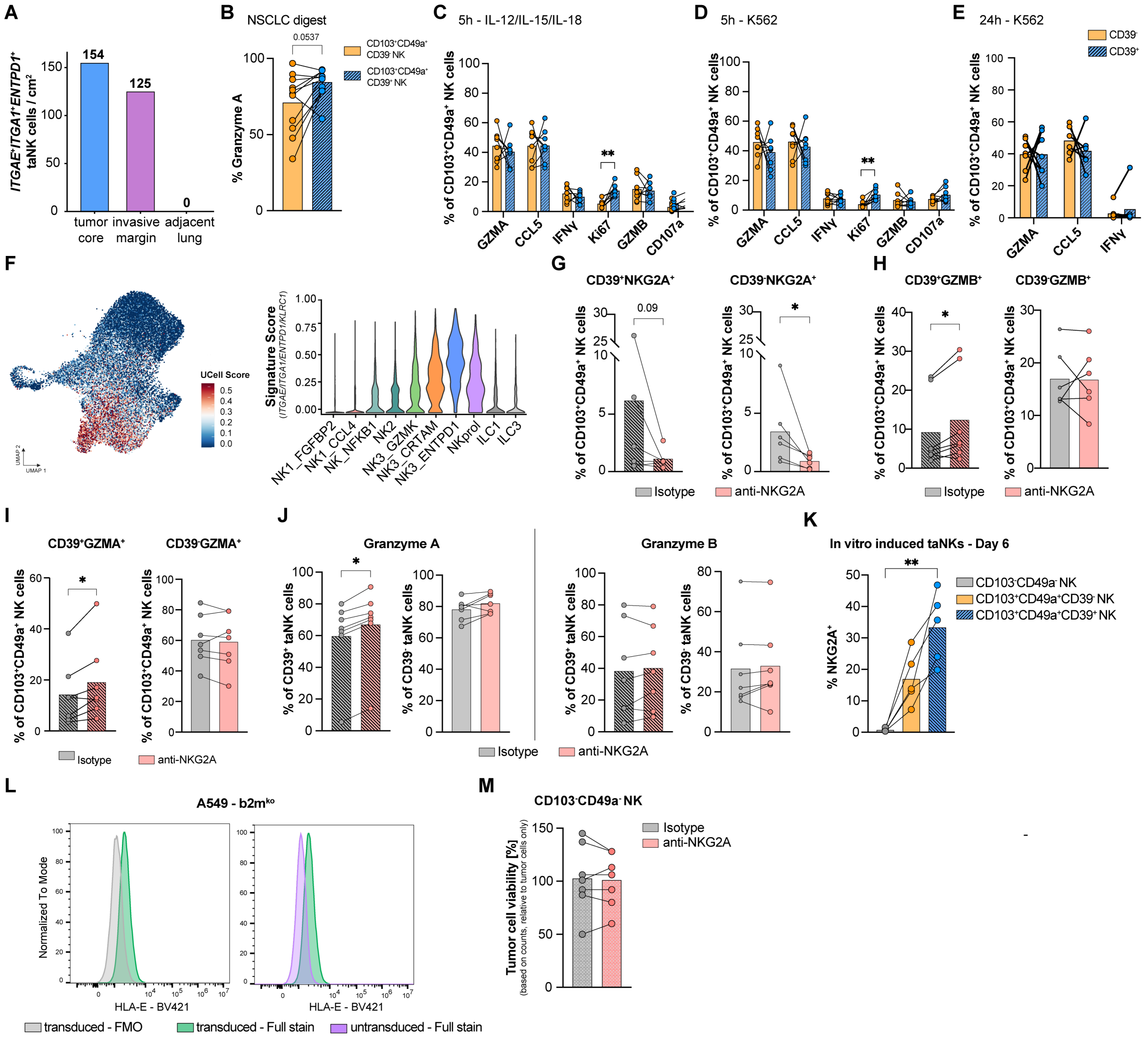

A
ITGAE+ITGA1+ENTPD1+ taNK cells / cm2
154
125
0
tumor core
invasive margin
adjacent lung
B
NSCLC digest
0.0537
% Granzyme A
CD103+CD49a+ CD39- NK
CD103+CD49a+ CD39+ NK
C
5h - IL-12/IL-15/IL-18
% of CD103+CD49a+ NK cells
GZMA
CCL5
IFNγ
Ki67
GZMB
CD107a
D
5h - K562
E
24h - K562
CD39-
CD39+
F
UCell Score
Signature Score (ITGAE/ITGA1/ENTPD1/KLRC1)
NK1_FGFBP2
NK1_CCL4
NK_NFKB1
NK2
NK3_GZMK
NK3_CRTAM
NK3_ENTPD1
NKprol
ILC1
ILC3
UMAP 1
UMAP 2
G
CD39+NKG2A+
CD39-NKG2A+
0.09
Isotype
anti-NKG2A
H
CD39+GZMB+
CD39-GZMB+
I
CD39+GZMA+
CD39-GZMA+
J
Granzyme A
Granzyme B
% of CD39+ taNK cells
% of CD39- taNK cells
K
In vitro induced taNKs - Day 6
% NKG2A+
CD103-CD49a- NK
CD103+CD49a+CD39- NK
CD103+CD49a+CD39+ NK
L
A549 - b2mko
Normalized To Mode
HLA-E - BV421
transduced - FMO
transduced - Full stain
untransduced - Full stain
M
CD103-CD49a- NK
Tumor cell viability [%] (based on counts, relative to tumor cells only)

**Fig. S12: Phenotypic and functional characterization of CD39$^{+}$ and CD39$^{-}$ taNK cell subsets. (A)** Quantitative density of taNK cells from 10X Xenium dataset co-expressing *ITGAE*, *ITGA1*, and *ENTPD1* across the indicated tissue regions (n = 1). **(B)** Barplot quantifying GZMA expression in CD103$^{+}$CD49a$^{+}$CD39$^{+}$ (dashed blue) compared to CD103$^{+}$CD49a$^{+}$CD39$^{-}$ (orange) NK cells from lung tumor digests (n = 11). **(C** to **E)** Frequencies of effector and proliferation markers in CD39$^{+}$ versus CD39$^{-}$ subsets of CD103$^{+}$CD49a$^{+}$ NK cells from NSCLC digests after stimulation with (C) IL-12/IL-15/IL-18 for 5 hours, (D) K562 cells for 5 hours, or (E) K562 cells for 24 hours (n = 7. **(F)** UMAP visualization and violin plots displaying signature scores for tissue-residency (*ITGAE, ITGA1, ENTPD1)* and checkpoint (*KLRC1* (NKG2A)) transcripts across NK cell clusters from NSCLC digests. **(G** to **J)** Percentages of the indicated CD103$^{+}$CD49a$^{+}$ NK cell populations in NSCLC-derived PDTFs following 48 hours treatment with isotype (grey) or anti-NKG2A (pink) (n = 7). **(K)** Quantification of NKG2A expression in CD103$^{-}$CD49$^{-}$, CD103$^{+}$CD49$^{+}$CD39$^{-,}$ CD103$^{+}$CD49$^{+}$CD39$^{+}$ on day 6 of the in vitro taNK cell platform. **(L)** Histograms validating HLA-E expression in A549-b2m$^{ko}$ cells transduced with HLA-E/b2m lentivirus, comparing transduced FMO (grey), transduced full stain (green), and untransduced full stain (purple). **(M)** Quantification of tumor cell viability (A549-b2mko-HLA-E) following co-culture with sorted CD103$^{-}$CD49$^{-}$ from the in vitro taNK platform treated with isotype or anti-NKG2A. Tumor cell viability is illustrated as a percentage relative to the tumor cells only control (n = 6) Significance was determined using [(B), (C), (D), (E), (G), (H), (I), (J) and (M)] paired Wilcoxon signed-rank test and (K) Friedmann test with post-hoc Dunn's multiple comparison. **P < 0.05, **P < 0.01, ***P < 0.001, ****P < 0.0001*)

### SUPPLEMENTARY TABLES

**Table S1: NSCLC patient characteristics used for single cell experiment.**

| Patient number | Sex | Year of birth | Smoking status | TNM | Stage | Histology |
|---|---|---|---|---|---|---|
| BS-1198 | female | 1955 | 30 py | pT2a pN0 | I | LUAD |
| BS-824 | male | 1941 | 12 py | pT1c pN1 | II | LUAD |
| BS-889 | female | 1962 | non-smoker | pT1 pN1 | II | LUAD |
| BS-897 | female | 1949 | 60 py | pT1b pN0 | I | LUAD |
| BS-1175 | female | 1955 | 14 py | pT2b pN2 | III | LUAD |
| BS-840 | male | 1945 | 40 py | pT3 pN1 | III | LUSC |
| BS-1308 | female | 1974 | non-smoker | pT3 pN0 | II | LUAD |
| BS-1314 | female | 1943 | 25 py | pT3 pN0 | II | LUSC |
| BS-1319 | male | 1962 | 50 py | pT3 pN1 | III | LUSC |
| BS-1322 | male | 1943 | 10 py | pT4 pN0 | III | LUSC |
| BS-1318 | female | 1962 | 45 py | pT1c pN0 | I | LUAD |

**Table S2: NSCLC patient characteristics used for functional and 10X Xenium application.**

| Patient number | Sex | Year of birth | Smoking status | TNM | Stage | Histology | Application |
|---|---|---|---|---|---|---|---|
| BS-719 | female | 1937 | 3 py | pT4 pN2 | III | LUSC | baseline staining |
| BS-1014 | female | 1075 | 45 py | pT4 pN1 | III | LUAD | baseline staining |
| BS-1100 | female | 1938 | 75 py | pT1 pN0 | I | LUAD | PDTF experiment |
| BS-1098 | male | 1954 | n/a | n/a | n/a | n/a | PDTF experiment |
| BS-1270 | female | 1962 | 40 py | pT4 pN1 | III | LUAD | activation assay |
| BS-1273 | male | 1949 | smoker | pT2a pN0 | I | LUSC | baseline staining |
| BS-1284 | male | 1954 | 50 py | pT1c pN2 | III | LUAD | Ex vivo Expansion |
| BS-1290 | male | 1947 | 60 py | pT2a pN0 | I | LUSC | baseline staining |
| BS-1305 | male | 1942 | smoker | pT4 pN0 | III | LUSC | baseline staining |
| BS-1324 | female | 1961 | 20 py | pT2a pN1 | II | LUAD | Ex vivo Expansion |
| BS-1340 | male | 1943 | n/a | pT2a pN0 | I | LUAD | baseline staining |

| | | | | | | | |
|---|---|---|---|---|---|---|---|
| BS-1341 | male | 1934 | 100 py | pT2 pN0 | I | LUAD | baseline staining, activation assay, Ex vivo Expansion |
| BS-1342 | male | 1957 | smoker | pT1 pN0 | I | LUSC | baseline staining |
| BS-1362 | male | 1958 | smoker | pT3 pN1 | III | LUAD | baseline staining, activation assay |
| BS-1371 | female | 1981 | non-smoker | pT1c pN0 | I | LUAD | baseline staining |
| BS-1373 | male | 1950 | 60 py | pT1b pN0 | I | LUSC | PDTF experiment |
| BS-1380 | male | 1947 | 30 py | pT1b pN0 | I | LUSC | baseline staining |
| BS-1383 | male | 1944 | 50 py | pT1a pN0 | I | LUAD | PDTF experiment |
| BS-1384 | male | 1954 | 75 py | pT1b pN0 | I | LUAD | PDTF experiment |
| BS-1387 | female | 1954 | 30 py | pT3 pN0 | II | LUSC | PDTF experiment |
| BS-1391 | female | 1949 | 60 py | pT4 pN2 | III | LUSC | baseline staining, PDTF experiment |
| BS-1400 | male | 1947 | 45 py | pT3 pN0 | II | LUSC | baseline staining, activation assay |
| BS-1405 | female | 1957 | 40 py | pT1c pN0 | I | LUAD | baseline staining |
| BS-1410 | female | 1962 | 40 py | pT2a pN0 | I | LUAD | activation assay |
| BS-1414 | male | 1946 | 20 py | pT1c pN0 | I | LUAD | activation assay, Ex vivo Expansion |
| BS-1422 | male | 1949 | 50 py | pT2a pN0 | I | LUAD | activation assay |
| BS-1437 | male | 1958 | 80 py | pT3 pN0 | II | LUAD | baseline staining, activation assay |
| BS-1474 | male | 1941 | 80 py | pT1c pN0 | I | LUAD | Ex vivo Expansion |
| BS-1476 | female | 1955 | 30 py | pT0 ypN0 | I | LUAD | Ex vivo Expansion |

| | | | | | | | |
|---|---|---|---|---|---|---|---|
| BS-1478 | female | 1955 | 50 py | pT2a pN0 | I | LUAD | Ex vivo Expansion |
| BS-1485 | male | 1962 | 30 py | ypT2a ypN2 | III | LUAD | Ex vivo Expansion |
| BS-1489 | female | 1944 | 25 py | pT2a pN1 | II | LUAD | Ex vivo Expansion |
| LuT-722 | male | 1951 | non-smoker | pT4, pN2 | III | LUAD | 10X Xenium |

**Table S3: Antibodies used for flow cytometry experiments.**

| Antigen | Fluoro-chrome | Clone | Cat# | Supplier | RRID | Concen-tration |
|---|---|---|---|---|---|---|
| CD45 | BV510 | HI30 | 304036 | Biolegend | AB_2561940 | 100ug/ml |
| CD45 | AF532 | HI30 | 58-0459-42 | eBioscience | AB_11218673 | 12ug/ml |
| CD45 | BV711 | HI30 | 304050 | Biolegend | AB_2563466 | 50ug/ml |
| CD14 | FITC | M5E2 | 301804 | Biolegend | AB_314186 | 400ug/ml |
| CD19 | FITC | HIB19 | 302206 | Biolegend | AB_314236 | 400ug/ml |
| CD3 | FITC | HIT3a | 300306 | Biolegend | AB_314042 | 200ug/ml |
| CD3 | AF647 | UCHT1 | 300416 | Biolegend | AB_389332 | 100ug/ml |
| CD8 | BV421 | SK7 | 344748 | Biolegend | AB_2629584 | ND (1:100) |
| CD8 | APC | SK1 | 344722 | Biolegend | AB_2075388 | 25ug/ml |
| CD56 | PE | 5.1H11 | 981202 | Biolegend | AB_2715758 | 80ug/ml |
| CD56 | BV785 | 5.1H11 | 362550 | Biolegend | AB_2566059 | 100ug/ml |
| CD103 | BUV661 | BER-ACT8 | 749993 | BD Biosciences | AB_2874215 | 0.2mg/ml |
| CD49a | BUV805 | SR84 | 749261 | BD Biosciences | AB_2873639 | 0.2mg/ml |
| CCL5 | PE | VL1 | 515504 | Biolegend | AB_2561328 | 25ug/ml |
| CD16 | BUV496 | 3G8 | 612944 | BD Biosciences | AB_2870224 | 400ug/ml |
| Granzyme A | PE-Cy7 | CB9 | 507221 | Biolegend | AB_2721667 | 4ug/ml |
| Granzyme B | AF700 | QA16A02 | 372222 | Biolegend | AB_2728389 | 25ug/ml |
| NKG2A | BV650 | 131411 | 747920 | BD Biosciences | AB_2872381 | 0.2mg/ml |
| NKG2C | BV711 | 134591 | 748164 | BD Biosciences | AB_2872625 | 0.2mg/ml |
| CD39 | BV605 | A1 | 328236 | Biolegend | AB_2750430 | 100ug/ml |
| CTLA4 | PE-Cy5 | BNI3 | 555854 | BD Biosciences | AB_396177 | ND (1:100) |

| | | | | | | |
|---|---|---|---|---|---|---|
| IFN-ɣ | BV421 | 4S.B3 | 502532 | Biolegend | AB_2561398 | 100ug/ml |
| CD107a | APC-H7 | H4A3 | 561343 | BD Biosciences | AB_10644020 | 50ug/ml |
| Ki67 | APC | Ki-67 | 350514 | Biolegend | AB_10959327 | 50ug/ml |
| CD73 | BV421 | AD2 | 344008 | Biolegend | AB_11204424 | 100ug/ml |

ND = unknown
